# Self-organized hemanoids derived from human iPSCs create a niche that produces definitive extraembryonic hematopoiesis

**DOI:** 10.64898/2026.05.05.722134

**Authors:** Afrim Avdili, Martina Auer, Dagmar Brislinger, Dagmar Kolb, Gerit Moser, Andreas Reinisch, Gerald Hoefler, Claudia Bernecker, Julia Fuchs, Julia Feichtinger, Peter Schlenke, Isabel Dorn

## Abstract

Manufacturing red blood cells (RBCs) from human induced pluripotent stem cells (iPSCs) can improve our understanding of embryonic erythropoiesis, foster innovative treatments for RBC-related diseases, and ultimately address clinical blood supply shortages. However, existing systems face low efficiency, enucleation failure, and uncertainty about the developmental wave of cultured RBCs. We successfully used self-organized hemanoids to improve iPSC-derived RBC generation. Based on the hypothesis that cellular interactions and 3D organization promote hematopoietic cell fate, we aimed to thoroughly characterize hemanoids. We visualized the spatiotemporal emergence of hematopoiesis by generating a CD43-GFP reporter iPSC line. Imaging and spatial transcriptomics analysis provided detailed insight into the hemanoid architecture, identifying stromal cells and hepatoblasts as potential erythropoiesis-supportive elements. The developmental stage mirrors extraembryonic hematopoiesis. Given the difficulties of accessing these early stages in vivo, our system offers a platform not only for further clinical translation but also for exploring human embryonic blood wave dynamics.

## Introduction

Culturing human RBCs ex vivo from induced pluripotent stem cells (iPSCs) has enormous potential to expand diagnostic and therapeutic options for RBC-related diseases and to address the increasing shortage in clinical blood supply.^1,2^ Furthermore, it offers a unique opportunity to deepen our limited understanding of human developmental erythropoiesis. Early developmental stages are not readily accessible in humans, and animal models, like mice, differ in key aspects.

The first blood cells originate in the yolk sac (YS), supplying the developing embryo with nutritional, metabolic, and oxygen support. Within the first 18 days post-conception (dpc) (Carnegie stage (CS) 7-8), mesenchymal cells adjacent to the endoderm differentiate into hematopoietic cells (HC) and become enveloped by endothelial cells, which later form the YS vascular plexus.^3–5^ This first wave of “extraembryonic primitive hematopoiesis“ generates RBCs, megakaryocytes, and macrophages. The YS is also the origin of the second wave, termed “extraembryonic definitive hematopoiesis”, generating erythro-myeloid progenitors (EMPs) from the endothelium through a gradual process known as endothelial-to-hematopoietic transition (EHT) (∼28-35 dpc, CS 13-15). With the onset of fetal circulation, EMPs leave primitive blood islands and migrate from the YS to the FL, where they produce a broader spectrum of HCs, including granulocytes, monocytes, and mast cells. Both extraembryonic waves are ultimately replaced by a third wave of intraembryonic hematopoiesis originating in the aorto-gonad-mesonephron (AGM) region (∼30-32 dpc, CS14). AGM-derived hematopoietic stem cells (HSCs), possessing self-renewing properties and enhanced lymphoid potential, initially colonize the FL alongside EMP-derived cells, before migrating into the bone marrow (BM) to sustain lifelong hematopoiesis.^5–7^ The different waves of human hematopoiesis overlap temporally and spatially throughout development. Preliminary exploration of cellular morphology and cell surface marker expression has yielded no clear markers for assigning individual cells to a specific wave. A distinction is likely possible based on differences in gene expression profiles, such as HSC-specific signature genes or globin genes.^8–13^ Primitive RBCs are characterized by expression of embryonic hemoglobins (embHb) Gower I (ζ_2_ε_2_) and Gower II (α_2_ε_2_). While EMP- and HSC-derived erythropoiesis in the FL synthesize predominantly fetal hemoglobin (HbF, α_2_γ_2_), BM-derived cells produce adult hemoglobin (HbA, α_2_β_2_).^4^

Since the discovery of iPSCs, several culture systems have been developed to model human erythropoiesis. To simulate the complex in vivo situation, researchers use extensive cytokine stimulation in combination with digestion and purification steps. Despite significant progress in recent years, established systems remain severely limited by low cellular yields and a failure to achieve terminal enucleation (below 10%).^2,14,15^ In addition, it remains unclear which developmental wave iPSC-derived cultured RBCs (cRBCs) represent. Because cellular interactions are essential for cell fate decisions, organoid-based systems have been established to differentiate pluripotent cells more effectively into various cell types.^16–19^ As the environment may also substantially impact hematopoietic development ^20,21^, our group developed a simplified erythropoiesis model based on the formation and maintenance of self-organized 3D complexes, termed „hemanoids“.^22,23^ Minimal cytokine stimulation (inter-leukin-3 (IL-3), stem cell factor (SCF), and erythropoietin (EPO))^24–27^ results in the continuous release of HCs from hemanoids into the culture supernatant over several weeks, which can be further differentiated into RBCs. High expansion and enucleation rates (up to 60%) already enabled us to confirm that cRBCs exhibit morphological, biomechanical, and blood-group antigen expression profiles comparable to those of cord blood-derived reticulocytes.^22,23^ Interestingly, by altering cytokine stimulation, the system can produce macrophages and granulocytes and is further scalable in stirred bioreactors.^28,29^

Despite these promising results, the underlying erythropoiesis supporting mechanisms remain unclear. For further improvement and meaningful future application of the system, it is crucial to comprehend the tissue architecture of hemanoids, the spatiotemporal development of hematopoiesis inside hemanoids, and to clarify which developmental wave the generated erythropoiesis corresponds to. To achieve these objectives, here we generated a CD43-GFP reporter iPSC line and tracked hematopoietic emergence within hemanoids. We investigated the hemanoids architecture by immunohistochemistry (IHC) and scanning transmission electron microscopy (STEM). Additionally, we performed spatial transcriptomics (ST) analysis to correlate histological characteristics with the transcriptional profile. We identified stromal elements and hepatoblasts as potential hematopoiesis-supportive interaction partners. Results argue for a developmental stage of EMP-derived hematopoiesis. The hemanoid model thus offers a unique platform for modeling definitive extraembryonic erythropoiesis spanning the undifferentiated iPSC stage through the enucleated RBC stage. Because in vivo access to these stages is limited, our model bridges existing gaps and offers valuable insights into embryonic erythropoiesis, potentially serving as a platform for future clinical applications.

## Results

### Hemanoids enable hematopoietic and erythroid differentiation of human iPSCs

We initially applied three iPSC lines^30–32^ as biological replicates to our culture system (**Figure 1**).^22,23^ After about five days of cytokine stimulation (phase I), self-organized three-dimensional hemanoids were established, consisting of an adhesive stromal layer, spherical structures, cell-dense areas, and, in individual cases, macroscopically detectable red islands. Throughout culturing, the size of the hemanoids increased significantly (**Figure S1A**). From approximately day 14 onwards, hemanoids continuously released CD43+ HCs (purity > 95% measured by flow cytometry) into the supernatant (**Figure S1B**), reaching a cumulative number of 3.1×10^6^ ± 1.7×10^6^ CD43+ cells over 5 weeks (n=6, mean ± SD; per one well of a six-well plate containing 1-2 hemanoids). Thereafter, the potential of hemanoids to release HCs was exhausted. For further erythroid differentiation, released CD43+ cells were repeatedly collected and subjected to RBC differentiation^33^ over an additional 18 days (phase II). Cells showed homogenous maturation into 99% GPA+/Band3+ erythroid cells^34^ (**Figure S1C**). On day 18, all cells expressed hemoglobin, and the percentage of enucleated cells exhibiting the typical RBC morphology reached 39.1% ± 16.4% (**Figures S1D-E**). A mean cumulative expansion of 3,673 ± 2,037-fold was observed during the 18-day erythroid differentiation phase (**Figure S1F**). These results are comparable to those reported in our previous study^22^, despite being derived from different iPSC sources. They, therefore, confirm the robustness and reproducibility of the hemanoid system. The advantages of this system are i) minimal handling, ii) low cytokine supplementation (only SCF, EPO, IL-3), iii) the continuous release of a pure population of CD43+ HCs into the supernatant, and iv) enhanced expansion and enucleation of erythroid cells. We hypothesize that these benefits arise from a specialized microenvironment and cellular interactions within the hemanoids.

**Figure 1.**
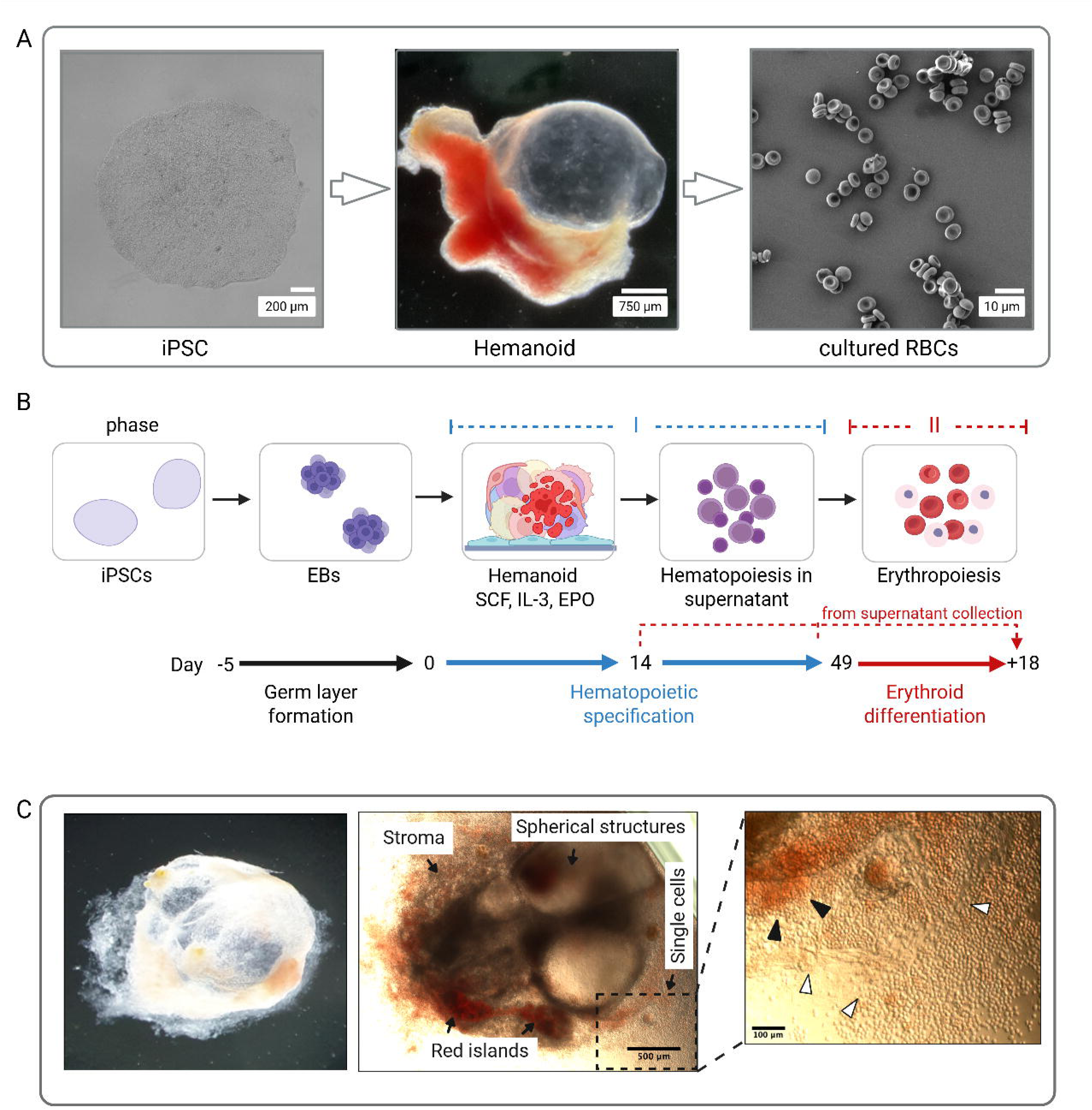
Hemanoid formation and erythroid differentiation of human iPSCs. (see also Figure S1). (**A)** Erythroid differentiation through 3D-hemanoids. Left: Undifferentiated iPSCs; middle: self-organized Hemanoid; right: Electron microscopy image of hemanoid-derived cultured RBCs (scale bars: 200 µm, 750 µm, 10 µm). (**B**) Illustration of the cell culture workflow. Days -5 to 0: Induction of germ layer formation by embryoid body (EB) formation. **Phase I** (days 0 – up to day 49): Hematopoietic specification / hemanoid formation in APEL™ medium supplemented with EPO, SCF, and IL-3. Starting around day 14, hemanoids continuously released CD43+ HCs into the supernatant. **Phase II** (+ 18 days): Erythroid differentiation of cells harvested between days 14 and 49 from the hemanoid supernatant (dashed red line) over an additional 18 days. (**C**) Left and middle: Representative brightfield images of a three-week-old hemanoid generated from PEB-AL#6 iPSCs, showing spherical structures, cell-dense areas with red islands, and an adherent stromal cell layer (Primovert Zeiss, 4x, scale bar: 500 µm). Right: Magnification of the rectangular area, representing red islands (black arrow), parts of the stromal layer (white arrow), covered by single cells released into the supernatant (scale bar: 100 µm).

### Spatiotemporal emergence of CD43+ hematopoietic cells

Because CD43 (leukosialin) is considered the first specific pan-hematopoietic cell-surface marker during human pluripotent stem cell differentiation^35^, we generated a CD43-GFP fluorescent reporter iPSC line (CD43R-iPSC) to monitor the emergence and subsequent distribution of HCs within hemanoids. Using CRISPR/Cas9 gene editing, we targeted the AAVS1 safe harbor locus on chromosome 19 to introduce the CD43_GFP vector construct through adeno-associated virus serotype 6 (AAV6)-mediated homology-directed repair (HDR)^36–39^, as shown in **Figure S2**. Successfully manipulated cells from the iPSC lines PEB-AL#6^30^ and CM1#1^32^ were clonally expanded. Precise CD43 reporter knock-in into the AAVS1 locus was confirmed by PCR genotyping (**Figure S2E**). The CD43R-iPSCs showed comparable hematopoietic and erythroid potential as their parental cell lines, as demonstrated by the number of CD43+ cells released into the supernatant, hematopoietic colony-forming potential in semisolid media, and erythroid differentiation capacity (**Figure S3**).

To identify the earliest CD43-GFP+ cells within hemanoids, we conducted time-lapse microscopy starting on day 5 of hematopoietic specification (phase I) (**Figure 2A**). The first GFP-expressing cells emerged on day 6, as confirmed by three independent experiments. CD43-GFP+ cells initially appeared in specific locations within cell-dense areas and subsequently expanded from there for approximately 5 weeks. GFP+ cells migrated within hemanoids, before being released as single cells into the supernatant (**Figures 2B-D)**. Flow cytometry characterization following cell-surface CD43 staining confirmed the correlation between endogenous CD43 gene expression and GFP expression from the inserted CD43-reporter construct (**Figures 2E and 2F**). Before their release, GFP+ HCs strongly adhered to the hemanoid surrounding stromal layer (**Figure 2G**). This adhesion was resistant to disruption, e.g., by extensive washing steps.

**Figure 2.**
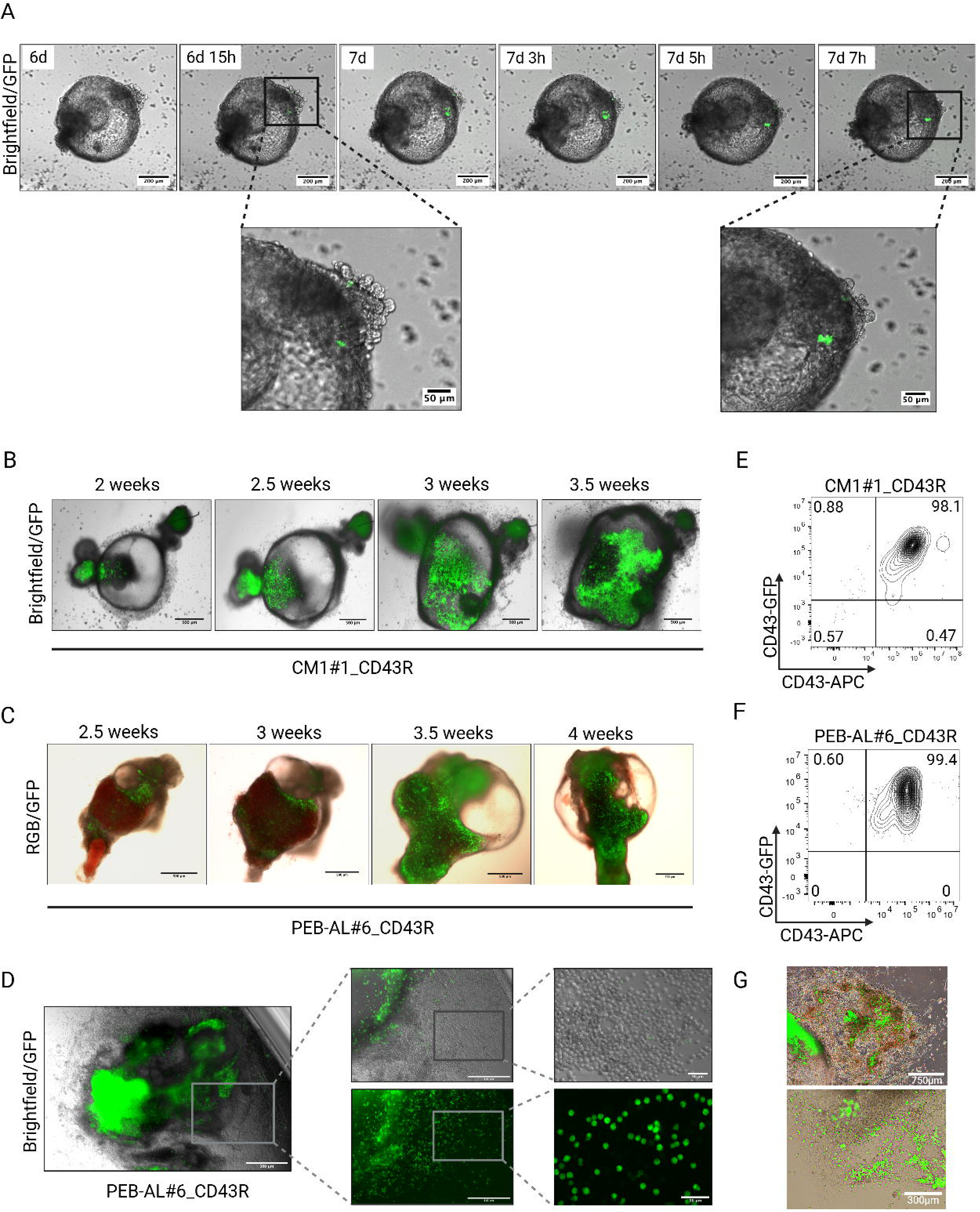
Emerging hematopoiesis inside hemanoids generated from CD43-GFP re-porter iPSC lines. (see also Figures S2 and S3). (**A**) Representative time-lapse microscopy images of a PEB-AL#6_CD43R iPSC-derived hemanoid expressing CD43 tagged with GFP. After 6 days and 15 hours in culture, the first GFP+ cells were detected (scale bar: 200 µm, magnification 50 µm). (**B and C**) Representative fluorescence microscopy images obtained between weeks 2 and 4 of a hemanoid from CM1#1_CD43R iPSCs and PEB-AL#6_CD43R iPSCs (scale bar: 500 µm). Red hemoglobin-positive areas are CD43-negative, consistent with CD43 downregulation during terminal RBC maturation. (**D**) Release of GFP+ single cells from a PEB-AL#6_CD43R hemanoid into the supernatant. Shown are microscopic magnifications of the rectangular areas (scale bars: 500 µm, 300 µm, and 50 µm). (**E and F**) Single cells released from CM1#1_CD43R and PEB-AL#6_CD43R hemanoids were stained against CD43 (APC) to confirm endogenous CD43 expression of GFP+ cells. (**G**) 4-weeks old hemanoids derived from PEB-AL#6_CD43R_iPSCs (top) and CM1#1_CD43R_iPSCs (bottom). GFP+ HCs demonstrate close contact with the stromal cell layer (scale bars: 750 µm and 300 µm).

### The heterogeneous tissue organization of the hemanoid reveals its complexity

Hemanoids consistently form blister-like structures, often more than one, filled with fluid (**Figure 3A**). In addition, they develop a stromal cell layer that extends beyond the complex and mediates adhesion of the hemanoid to the culture plate (**Figure 3B**). In our previous study^22^, we observed that both features are crucial for successful hematopoietic induction. In contrast to the pure HC population released into the supernatant, we found only 49.6% ± 8.8% CD43+ HCs within the hemanoids, measured by flow cytometry after enzymatic digestion (**Figure 3C**). Hematoxylin/eosin (HE)-stained tissue sections obtained between 16 and 28 days of hematopoietic specification (phase I) showed morphological features comparable to those of ectodermal (neural crest tube-like structures), mesodermal (stromal tissue and blood cells), and endodermal origin (glandular structures) (**Figures 3D and S4A**). Especially at early stages (<20 days), CD43+ HCs were primarily found within VE-Cadherin+ (CD144) vessel-like structures that resembled YS blood islands morphologically. Vessels were surrounded by vimentin+/β-laminin+ mesenchymal-like cells (**Figures 3E and S4B-C**). CD163+ macrophages were the only HC type predominantly located outside blood vessels and within this stromal compartment (**Figure S4D**). A notable difference in older hemanoids (day 28) was the predominance of hematopoiesis outside blood vessels and within the mesenchymal compartment (**Figures 3F and S4B**). The hematopoietic compartment itself contained erythroid precursor cells, granulocytes, monocytes, megakaryocytes, platelets, and mast cells (**Figures 3G-H**) as confirmed by antibody-mediated staining (**Figures S4D-H**). CD61+ megakaryocytes exhibited a small cell size (∼20 µm) typical of their juvenile maturation stage^40^ (**Figure S4E**). A striking feature was the predominance of eosinophilic granulocytes (**Figures 3G and S4F**). Although all HC types were detectable in day 16 hemanoids, mast cells and granulocytes became more abundant in older hemanoids.

**Figure 3.**
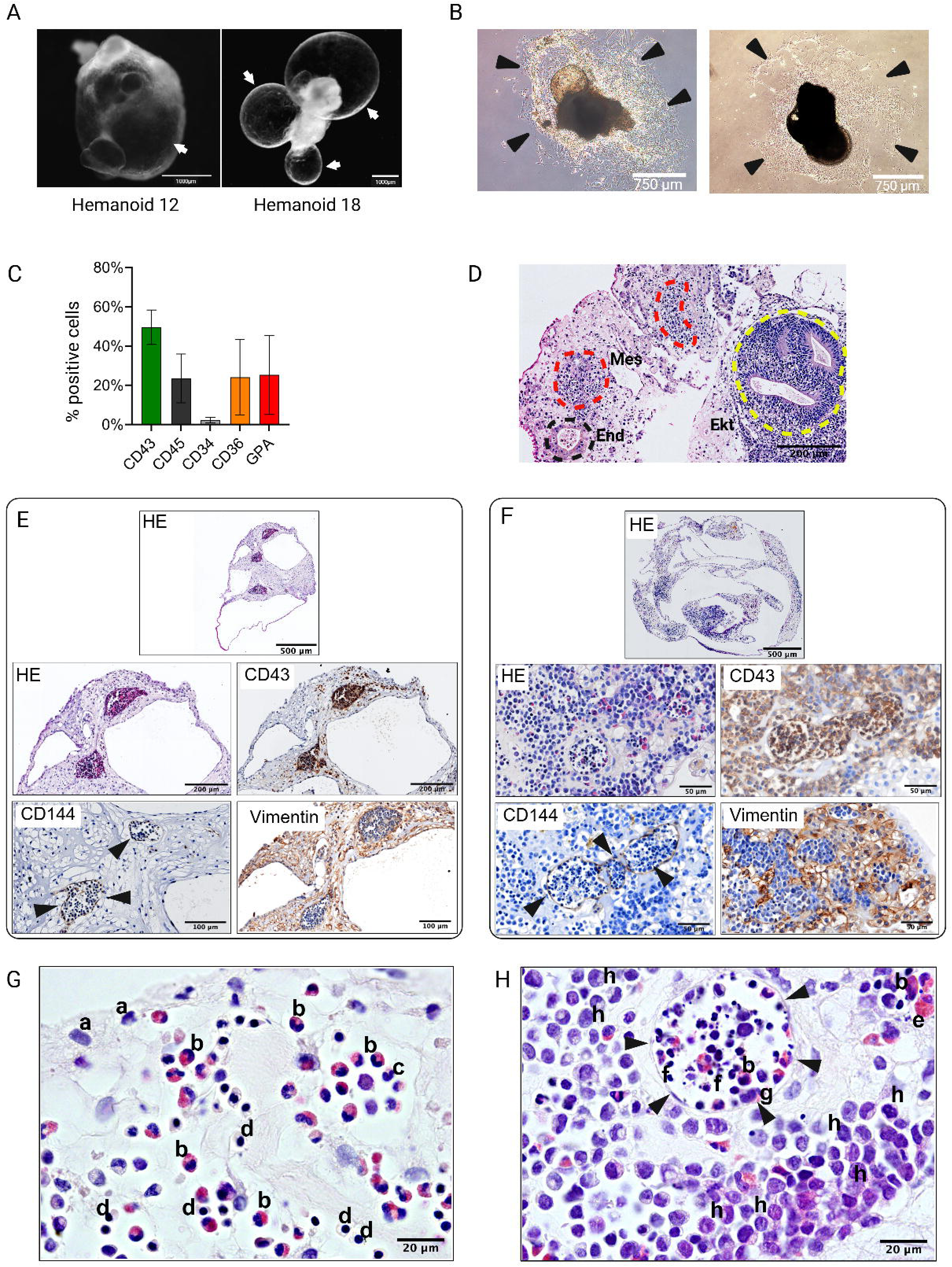
Immunohistochemistry-based characterization of hemanoids. (**A**) Representative images of two different hemanoids. White arrows indicate blistered, fluid-filled are-as (scale bar: 1mm). (**B**) Brightfield images of two PEB-AL#6 hemanoids (3 and 4.5 weeks). Black arrows indicate the stroma layer that attaches them to the tissue culture plate (scale bar: 750 µm). (**C**) Percentage of cells stained positive for hematopoietic cell surface markers after enzymatic dissociation of hemanoids, and analyzed by flow cytometry (n=6; 3 biological replicates; mean ± SD). (**D**) HE-stained hemanoid section showing morphology of mesodermal (Mes) tissue with hematopoietic areas (red dashed line), ectodermal structures (Ect) (yellow dashed line), and endodermal glandular structures (End, black dashed line) (scale bar: 200 µm). (**E and F**) HE-stained tissue sections of two different hemanoids. Shown are cross-sections and magnifications, co-stained by horseradish peroxidase for CD43 (hematopoiesis), CD144 (VE-cadherin, endothelial cells), and vimentin (mesenchymal cells). In (E), CD43+ HCs are found within CD144+ vessel-like structures (black arrows), surrounded by vimentin+ mesenchymal-like cells. In (F), CD43+ HCs are found both inside (black arrows) and outside the CD144+ vessel, within a network of vimentin+ cells. (**G and H)** Brightfield images of HE-stained hemanoids (100X oil, scale bar: 20 µm), showing hematopoietic areas containing erythroid cells, myeloid cells, and thrombocytes. (G) HCs within the stromal network, with a predominance of eosinophils. (H) small vessel (black arrows) containing, e.g., eosinophils and platelets, surrounded by myeloid cells. (**a** Mesenchymal cell; **b** Eosinophilic granulocyte; **c** Neutrophilic granulocyte; **d** Erythroid precursor cell; **e** Mast cell; **f** Platelets; **g** Monocyte; **h** Myeloid precursor). See **Figure S4** for confirmation of cell types by specific antibody staining.

To extend our imaging to the nanometre level, we performed STEM analysis of tissue sections (**Figures 4 and S5**). Ultrastructural analysis confirmed our IHC findings, showing blood islands with various HC types lined by endothelial cells in day 17 hemanoids (**Figures 4A-C and S5A**). The surrounding stromal compartment was characterized by a loose network of mesenchymal-like cells, fibroblasts, collagen fibers, and individual tissue macrophages (**Figure 4D**). By day 28, the hemanoids exhibited a significant transformation; the stromal tissue had become highly organized, with a dense network of collagen fibers produced by an increased number of stromal cells, yet hematopoiesis had declined (**Figure S5B**). Blood islands became less distinct and showed signs of disruption, while HCs were primarily located within the stroma (**Figure S5C**). Already by day 17, STEM revealed cell-cell interactions between erythroid precursors and other cells, suggesting metabolite exchange and mutual influence (**Figure S5D**). The outer surface of the hemanoid was covered by a conspicuous epithelial layer of endodermal origin (AFP+, ß-laminin+, HLA-G-, CK7-) (**Figure S5E**). This layer was further characterized by a basement membrane, intracellular vesicles, and cell extensions at the outer surface. Cells were partially connected via desmosomes.

**Figure 4.**
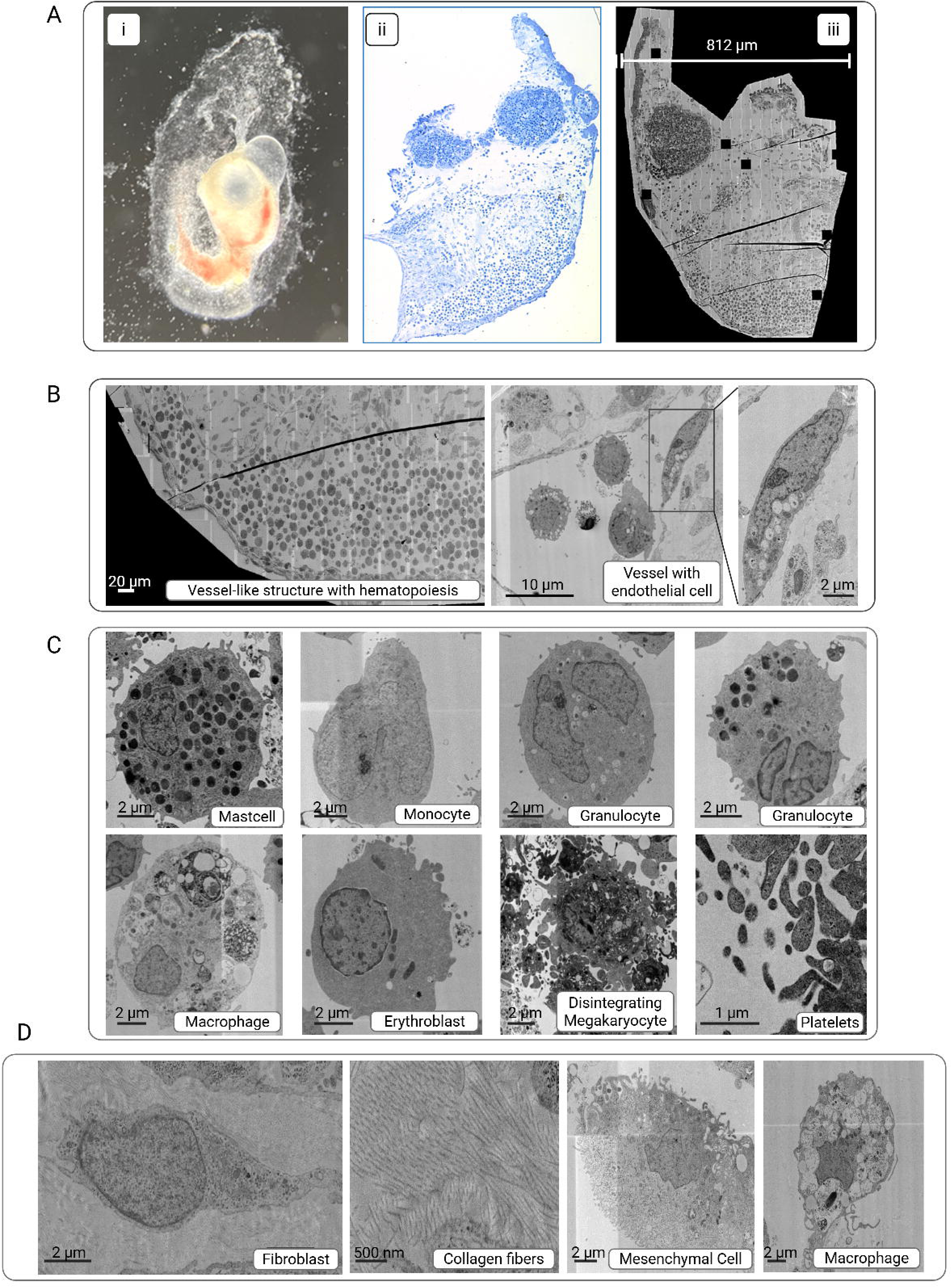
STEM analysis of a hemanoid (day 17) derived from PEB-AL#6. (see also Fig-ure S5). (**A**) The identical hemanoid is shown i) as an adherent hemanoid in the dish, ii) after fixation, sectioning, and toluidine blue staining, and iii) analyzed by STEM (diameter: 812 µm). (**B**) Magnifications from A, from left to right: Vessel containing HCs; HCs within the vessel; endothelial cell. (**C**) Magnifications from B: Different HC types inside vessels (scale bar: 2 µm, platelets 1µm). (**D**) Cells and fibers of the stromal compartment (scale bar: 2 µm, collagen fibers 500 nm).

### Spatial transcriptomics confirmed tissue across the three germ layers and distinct hematopoietic populations

To further extend our results to a molecular level and specify the developmental wave of hemanoid-derived hematopoiesis, we performed a 10X Visium spatial transcriptomics (ST) analysis of day 16 and day 28 hemanoids (**Figure 5A**). Day 16 was chosen based on previous results showing an initial increase in CD43+/CD45+ HCs at this culture day.^41^ Day 28 was selected as a later stage in hematopoietic development, prior to exhaustion of the system. Ten to eighteen hemanoids from PEB-AL#6 iPSCs were pooled to cover the capture area of the ST slide (6.5 mm x 6.5 mm, **Figure S6A**). We initially focused on evaluating the day 16 data. RNA expression data from 1,002 spots in the capture area were categorized into 10 clusters and annotated using Azimuth with reference datasets for human embryonic development^42^ and tissue-specific marker genes (**Figures 5B-C and S6A**). Erythroid-megakaryocyte progenitors (EryMK), erythroid precursors (Erythroid), myeloid precursors (Myeloid), and stromal cells (Stroma) represented mesoderm-derived tissue. Hepatoblast-like cells (Hepatoblasts), intestinal/bronchoalveolar epithelial-like cells (Epithelial), and a cluster enriched in endodermal gene expression (Endo) represented the endodermal tissue, while neuroprogenitors (Neuro) and photoreceptor cells (PRC) were categorized as ectodermal-derived clusters. **Figure 5D** shows the expression of two representative marker genes per annotated cluster, such as *NES* and *MAP6* in neuroprogenitors (ectoderm), *SERPINA1* and *FGB* in hepatoblasts (endoderm), and *HAND1/2* in stromal cells (mesoderm) (**Figure 5D**). One cluster could not be assigned (undefined). It likely contains transcripts from a mixed cell population, as also visible in the heat map (**Figure S6B**). The contribution of more than one cell type to individual clusters was expected, given the 55 µm spatial resolution (spot size) achievable with the 10X Visium system.

**Figure 5.**
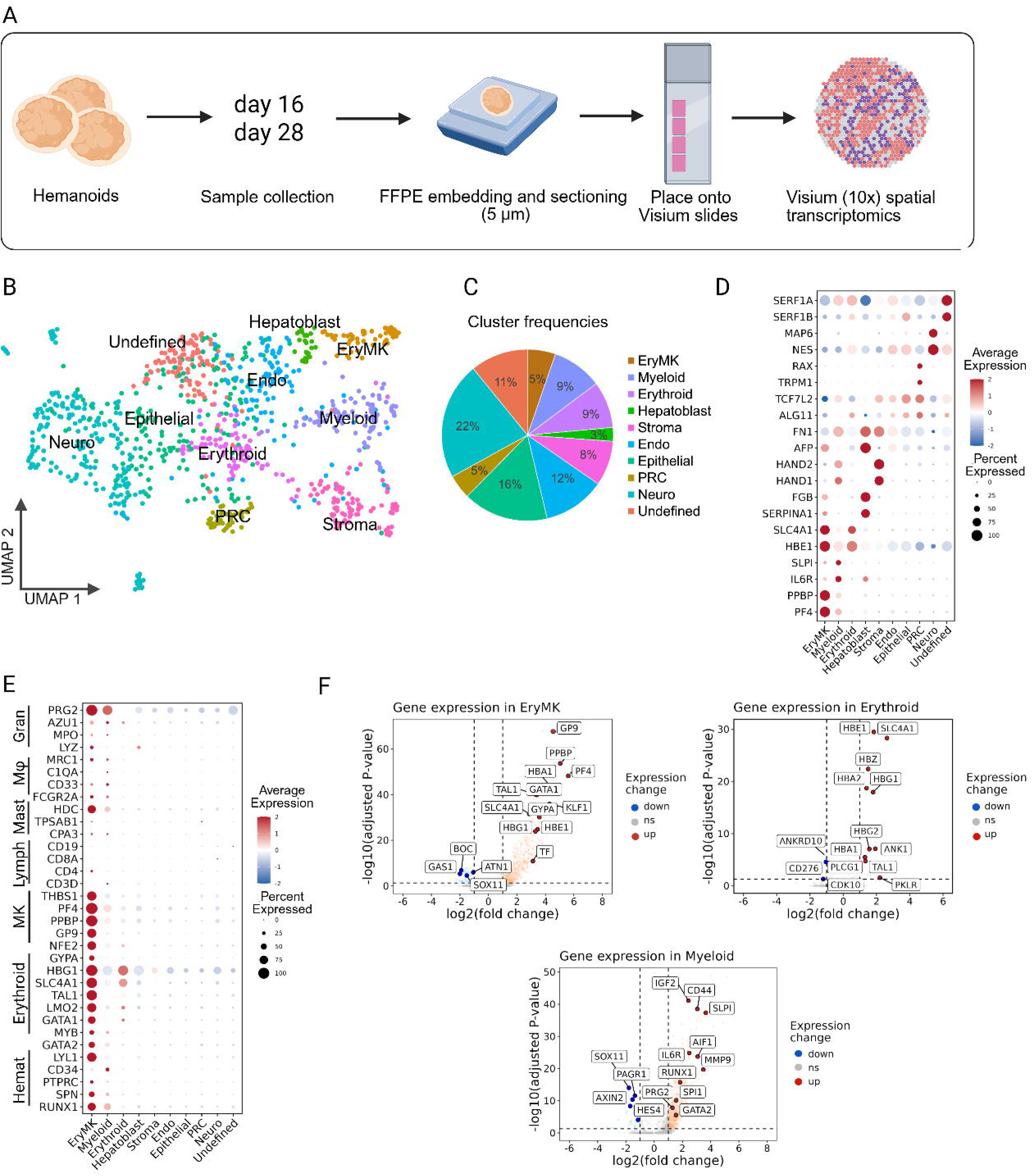
Spatial transcriptomics (ST) analysis and cluster annotation of day 16 hemanoids. (see also Figures S6-S8). (**A**) Schematic overview of the ST workflow. (**B**) Uniform Manifold Approximation and Projection (UMAP) visualization of ST-sequencing data. Colors indicate the 10 gene-expression-based clusters. (**C**) Percentage of spots contributing to each of the 10 identified clusters. (**D**) Dot plot showing the expression frequency (dot size) and the expression level (color intensity) of two marker genes in each cluster. (**E**) Dot plot showing the expression of canonical marker genes for the hematopoietic lineage (Gran (Granulocytes), Mϕ (Makrophages), Mast (Mast cells), Lymph (Lymphoid cells), MK (Megakaryocytes), Hemat (Hematopoietic cells). (**F**) Volcano plots showing upregulated and downregulated genes (log2-fold change) in the EryMK, Myeloid, and Erythroblast clusters compared to mean expression in all other clusters.

When all clusters were examined for a hematopoietic signature, this was mainly detectable in the EryMK cluster and, to a lesser extent, in the Myeloid and Erythroid clusters (**Figure 5E**). The EryMK and Myeloid clusters expressed common hematopoietic genes *RUNX1*, *SPN* (CD43), *and GATA2*. The EryMK cluster expressed genes relevant for platelet-mediated hemostasis, like *PF4* (platelet factor 4), *PPBP* (pro-platelet basic protein), and *GP9* (glycoprotein IX Platelet), but also erythroid-specific globin genes (*HBG1, HBE1*, *HBA1*), *GYPA* (glycophorin A), *SLC4A1* (Band3), and *GATA1* (**Figures 5E-F**). Upregulated genes within the Erythroid cluster encode essential RBC components, including globin genes (*HBG1*, *HBG2*, *HBE1*, *HBZ*, *HBA1/2*), *PKLR* (pyruvate kinase), *ANK1* (ankyrin), and *SLC4A1*. Based on the gene expression profile, the Erythroid cluster may encompass the more differentiated erythroid population, whereas the more immature progenitor population contributes to the EryMK cluster. The Myeloid cluster expressed the macrophage-associated genes *MRC1*, *CD33,* and *FCGR2A* (**Figure 5E**), as well as granulocyte-associated genes *PRG2, MMP9, IL6R, AIF1, and SLPI* (**Figure 5F**). Feature plots illustrating expression patterns for selected hematopoietic marker genes are shown in **Figure S7**. Gene set enrichment analysis (GSEA) revealed enrichment of the “response to oxygen” and “platelet activation” pathways in EryMK. Myeloid progenitors were enriched for “Leukocyte activation” and “Adaptive immune response” pathways (**Figure S8A**). Clusters were further validated using Gene Ontology (GO) analysis. In Enriched pathways in EryMK play a crucial role in erythrocyte differentiation and blood coagulation. In the Myeloid cluster, pathways were associated with the regulation of immune effector processes, leukocyte migration, and activation (**Figure S8B**).

#### Transcriptional profiles indicate a developmental stage that mirrors definitive extraembryonic hematopoiesis

Morphologically detectable red islands already indicate hemoglobin production within day 16 hemanoids. ST data confirmed the expression of globin genes in the Erythroid and the EryMK cluster (**Figures 6A-B**). Both clusters expressed *HBZ* and *HBA1/2* from the alpha-globin locus and embryonic *HBE1* and fetal *HBG1/HBG2* from the ß-globin locus, indicative of the synthesis of embHb (Gower I and Gower II) and HbF. Definitive *HBB* was barely detectable. Whereas primitive RBCs primarily express embHb, the coexpression of embHb and HbF is a typical pattern in EMP-derived erythropoiesis^4^ and was confirmed on a protein level by IHC staining of hemanoids (**Figure S9A**). Although hemanoids expressed *SOX6* and *BCL11A,* as repressors of embHb and HbF^43–45^, we did not observe their significant upregulation in one of the hematopoietic clusters (**Figures 6C and S9B-C**).

**Figure 6.**
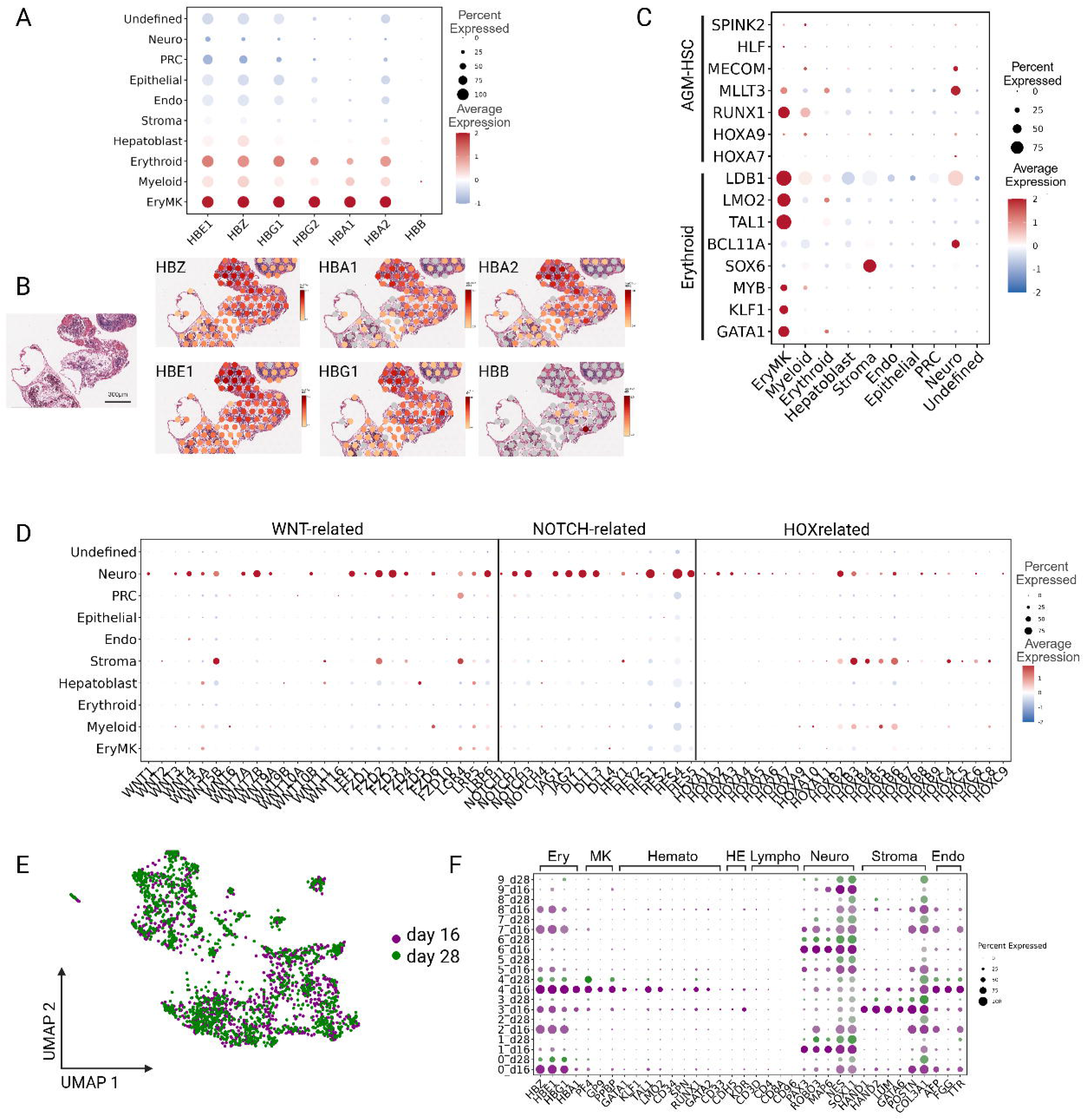
Developmental wave of hemanoid-derived hematopoiesis. (**A**) Dot plot showing the expression of genes encoding the alpha and beta chains of embryonic, fetal, and adult hemoglobin in day 16 hemanoids. (**B**) Globin gene expression overlay on Visium spots from a section of day 16 hemanoids (Loupe browser projection, log2 transformed UMI counts). (**C**) Expression of signature genes for AGM-derived HSCs^9^ and of core erythroid transcription factors^8^ in day 16 hemanoids (see also Figure S9). (**D**) Expression of genes related to WNT signaling, Notch signaling, and HOX gene clusters in day 16 hemanoids. (**E**) UMAP visualization of integrated ST data sets from day 16 and day 28 hemanoids, using Harmony (see Figures S10C and S11 for day 28 ST analysis). (**F**) Expression of cell-type-specific marker genes in the integrated day 16 (pink dots) and day 28 (green dots) data sets from (E): Erythroid (Ery), megakaryocytes (MK), hematopoietic (Hemato), hemogenic endothelium (HE), lymphoid (Lympho), neuro progenitors (Neuro), Stroma, endodermal (Endo). Dot size represents gene-expression frequency, and color intensity indicates expression levels.

*GATA1*, *KLF1*, *TAL1*, *LMO2*, and *LDB1*, as core erythroid TFs^46,47^, were expressed in the EryMK cluster (**Figure 6C**). Additional *MYB* expression confirms an advanced EMP stage, as *MYB* is not expressed in the primitive program.^48,49^ Moreover, we compared our dataset with a set of marker genes (*RUNX1, HOXA9, MLLT3, MECOM, HLF, SPINK2*), recently described by Calvanese et al.^9^, to distinguish intraembryonic AGM-derived HSCs from their extraembryonic progenitors. Although we observed expression of all six genes across the EryMK and the Myeloid cluster, there was no clear match of all markers to a single subcluster (**Figures 6C and S9B-C**). Enhanced lymphopoiesis as a hallmark of AGM-derived hematopoiesis was not detectable. Hematopoietic clusters showed minor expression of *CD3* (T-lymphocytes), but lacked expression for *MS4A1* (CD20) or *CD19* (B-lymphocytes) (**Figures 5E and S7**). IHC confirmed individual CD3+ cells, whereas CD20+ B cells were absent (**Figure S9D**). Therefore, we could not confirm a cell population comparable to AGM-derived hematopoiesis. The development of AGM-derived hematopoiesis is significantly influenced by WNT signaling (controlling early mesodermal patterning)^50^, NOTCH signaling (essential for arterial hemato-vascular development)^51,52^, and HOX pathways.^9,12^ Genes involved in WNT signaling, NOTCH genes (*NOTCH 1–4*), NOTCH ligands (*JAG1/2, DLL1*, *3,* and *4*), and target genes (*HEY1/2*, and *HES1-5*) were not significantly expressed in hematopoietic clusters. HOX gene expression was also sparse, although the EryMK and myeloid clusters indicate low *HOXA9* and *HOXA10* expression. In contrast to their low expression in the hematopoietic clusters, WNT, NOTCH, and HOX pathways were significantly upregulated in the Neuro cluster (**Figure 6D**).

### The transcriptional profile of hemanoids overlaps with that of human YS and FL

We set out to determine whether day 16 hemanoids show similarities with the human YS and FL, as sources of primitive and EMP-derived HCs. We integrated our ST data with two publicly available scRNA-seq datasets on the developing human YS at CS 10/11^53^ and 17^54^ using Harmony.^55^ After single-cell data processing and filtering, we retained gene expression data from 7,545 cells from CS 17 and 10,893 cells from CS 10/11. UMAP visualization reveals a similar expression pattern among certain clusters (**Figure S10A**). We further integrated our ST data with two datasets on human FL development: one from CS 20 and 23^53^, and the other from CD45+ isolated FL cells from post-conception weeks (wpc) 8–16.^11^ Our data also show overlap with cell populations during human FL development (**Figure S10B**).

### Day 28 hemanoids show a reduced hematopoietic potential and no further induction of AGM-derived definitive hematopoiesis

Since prolonged culturing of hemanoids might induce AGM-derived hematopoiesis, ST analysis was extended to day 28 hemanoids. RNA expression data derived from 1,349 spots of the capture area were categorized into 7 clusters and annotated using Azimuth with reference datasets for human embryonic development^42^ (**Figures S11A-B**). The mesodermal tissue was represented by an erythroid cluster (Erythroid), a cluster enriched for myeloid and megakaryocyte gene expression (Hemato), a stromal cluster (Stroma), and a separate myofibroblast cluster (Myofibroblast). Hepatoblast-like cells (Hepatoblast) and neuroprogenitors (Neuro) represented endodermal and ectodermal tissue. One cluster could not be assigned (Undefined). Interestingly, the hematopoietic compartment’s contribution to the overall gene expression profile decreased compared with day 16 (9% versus 23%), whereas the Neuro (37% versus 21%) and Stroma (21% vs 8%) contributions increased. **Figure S11C** shows the expression of three marker genes per cluster. A hematopoietic gene expression profile was primarily detected in the Hemato and Erythroid clusters (**Figure S11D**). Both expressed embryonic and fetal globin genes in the absence of ß-globin. In line with this, upregulation of *BCL11A* and *SOX6* by HCs was not detectable (**Figures S11E-F**). Investigation of hemato-poiesis-related signaling pathways NOTCH, WNT, and HOX, and expression of signature genes for AGM-derived HSCs^9^ gave no evidence of further induction of AGM-derived definitive hematopoiesis in day 28 hemanoids (**Figures S11F-G**). Transcriptome profiles of day 16 and 28 hemanoids were further integrated and batch-corrected using Harmony.^55^ Unsupervised UMAP clustering grouped the cell populations from the two samples into 9 clusters. Day 16 and day 28 samples displayed a comparable pattern of gene expression. However, the overall expression of hematopoiesis-related genes was reduced in day 28 samples. In particular, the expression of early hematopoietic markers CD34, SPN, RUNX1, and GATA2 was very low, whereas connective tissue genes like COL3A1 and neuroprogenitor-related genes like NES and SOX11 remained consistently expressed (**Figures 6E-F and S10C**).

### Stroma cells and hepatoblasts provide a supportive niche for early hematopoiesis

Microscopic evaluation of the ST tissue sections shows that already by day 16, hemanoids are capable of producing mature RBCs without undergoing the erythroid differentiation step (phase II) of our protocol (**Figure 7A**). This highlights the potential of hemanoids to create a supportive environment for erythroid development. In addition to the obvious interaction between HCs and stromal cells, ST analysis revealed a close proximity to hepatoblast-like cells. Cells with a hepatoblast-like gene expression profile were often located near primitive blood islands (**Figure 7B**). We analyzed gene expression patterns within these clusters, focusing specifically on genes encoding growth factors, adhesion molecules, and their counterparts.

**Figure 7.**
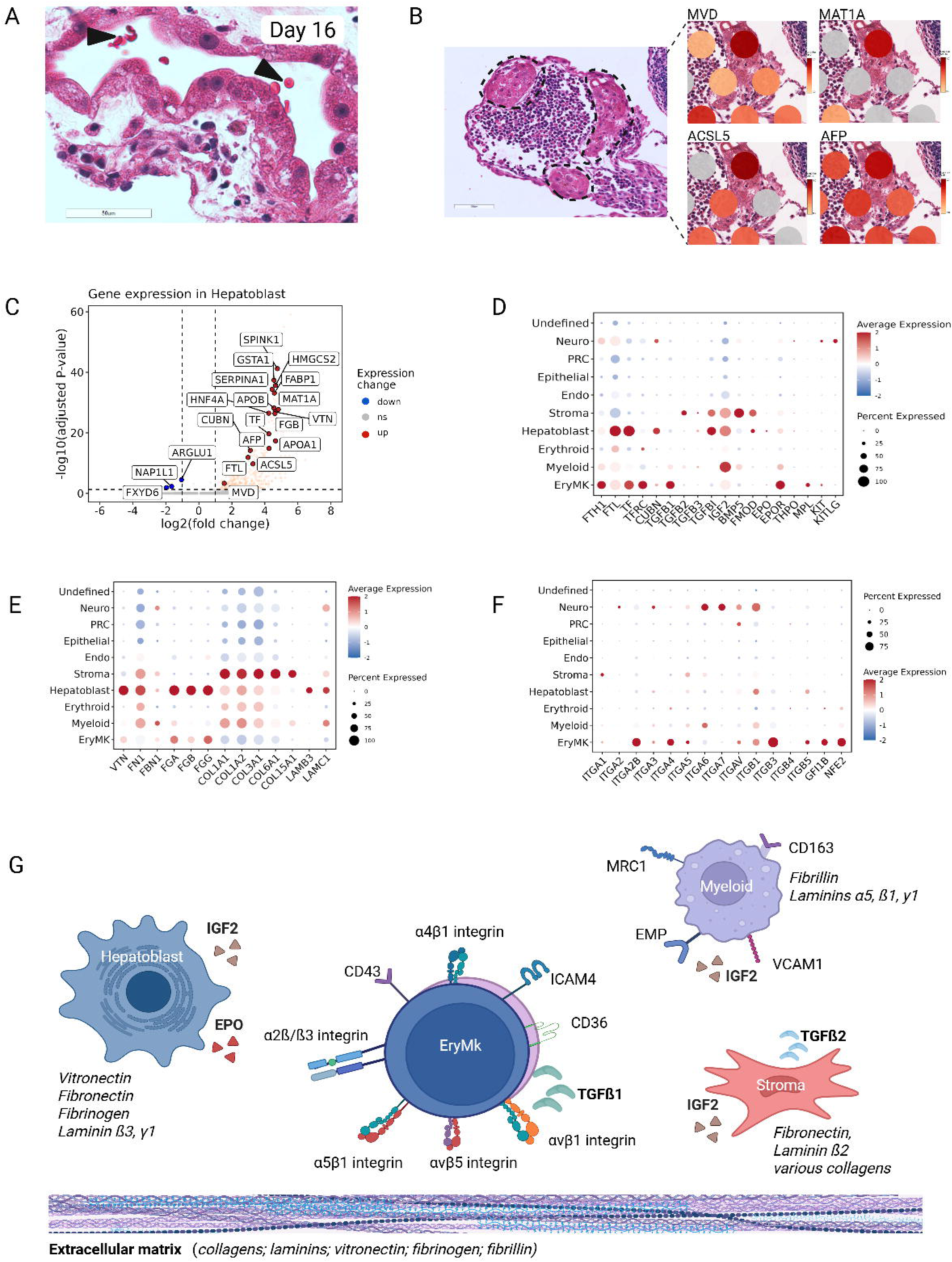
Cellular interactions of HCs (ST analysis of day 16 hemanoids) (see also Figures S12 and S13). (**A**) HE-stained hemanoid section on the ST slide showing terminal matured RBCs (black arrows) (scale bar: 50 µm). (**B**) Morphology of hepatoblasts (left, dotted lines) located near blood islands. Hepatoblast-specific marker gene expression overlay on Visium spots (Loupe browser projection). (**C**) Volcano plot showing upregulated and downregulated genes (log2 fold change) in the Hepatoblast cluster. (**D**) Dot plot showing the expression of genes encoding growth factors. (**E**) Expression of genes encoding ECM components. (**F**) Expression of genes encoding adhesion molecules. (**G**) Graphical illustration summarizing day 16 ST results regarding the expression of growth factors (bold), adhesion molecules, and ECM (Italic) in the EryMK, Myeloid, Hepatoblast, and the Stroma cluster.

The Hepatoblast cluster showed upregulated gene expression (e.g., *SERPINA1* (Alpha-1 Antitrypsin), *HNF4A* (Hepatocyte nuclear factor 4-alpha), and *ACSL5* (Acyl-CoA Synthetase)) and enrichment of pathways directly linked to liver cell metabolism (**Figures 7B-C and S12**). In addition to genes involved in lipid metabolism (e.g., *APOA, APOB, and ACSL5)*, expression of genes involved in iron and Vitamin B12 metabolism was observed, such as *FTL (*Ferritin-L chain, involved in iron storage, *TF* (transferrin), and *CUBN (*cubilin, a vitamin B12 receptor complex). The Hepatoblast cluster further expressed *IGF2* (insulin-like growth factor 2) and, to a minor extent, *EPO* (**Figures 7C-D and S12**). We found upregulation of genes encoding various ECM proteins, including vitronectin (*VTN*), fibronectin 1 (*FN1*), fibrinogens (*FGA, FGB, FGG*), and laminins β3 (*LAMB3*) and C1 (*LAMC1*) (**Figure 7E**). Recent reports on primary human YS hematopoiesis^10,56^, have identified an interaction between endodermal vitronectin and αvβ1-integrin, or the integrin subu-nits alpha2b, beta3, and beta5 on HCs, and between fibronectin and α4ß1- and αvß1-integrin on HCs. We confirmed upregulation of all the corresponding integrin-coding genes in the EryMK cluster (*ITGA2B, ITGA4, ITGA5, ITGAV, ITGB1, ITGB3, and ITGB5, as well as GFI1B and NFE2*, involved in *ITGB3* signaling^57^) (**Figure 7F**).

In the Stromal cluster, the transcriptional profile and GO enrichment analysis confirmed involvement in “Extracellular matrix organization” and “connective tissue development” (**Figure S13A-B**). Upregulated genes include those encoding various collagens, fibronectin (*FN1*), laminin-ß2 (*LAMB2*), and, to a lesser extent, fibrillin (*FBN1*) (**Figures 7E and S13C-E**). Several top upregulated genes regulate cellular development (*DSC3*, *GATA6*, *LMCD1), and* cardiac and angiogenic differentiation *(HAND1/2, TBX20, TNNT2, OLFML3).* We further observed increased activity in TGF-ß pathways, including *TGFB2*, *BMP5* (encoding a secreted TGFß ligand^58^), *FMOD* (coding fibromodulin, which regulates TGF-ß activity by sequestering TGF-ß in the ECM^59^), and *TGFBI* (TGF-ß-induced protein that influences cell adhesion). Notably, the EryMK cluster showed a significant rise in *TGFB1* expression (**Figure 7D, S13B and S13F**), while myeloid cells expressed *FBN1* (Fibrillin), which controls TGF-ß bioavailability and interacts with α5ß1- and αvß3-integrins (**Figure 7E**).^60,61^

In human BM and FL, erythroid maturation occurs in close contact with CD163+ macrophages in erythroblastic islands (EBI). Interaction is mediated by a4ß1-integrin, EMP, and ICAM4 on RBCs and VCAM1, EMP, and αvß3-integrin on macrophages.^62–65^ Although we found no obvious morphological correlates of EBIs, the myeloid cluster expressed *CD163* (confirmed by IHC staining), *MAEA* (coding EMP), *MRC1, VCAM1, and SIGLEC1,* in line with the phenotype of EBI macrophages. In addition to *ITGA4* and *ITGB1* (coding integrin α4β1-integrin), the EryMK cluster also expressed *ICAM4* as a potential interaction partner (**Figure S13G**).

## Discussion

Using an integrated imaging and transcriptomic approach, we obtained detailed insights into self-organized hemanoids that facilitate improved RBC generation from human iPSCs. The composition of the hemanoid is highly heterogeneous, comprising ectodermal, mesodermal, and endodermal tissue. The surface was covered by an epithelial layer of endodermal origin that may function in the exchange of nutrients, metabolites, and water between the hemanoid and the culture medium. Others reported trophoblast-like features.^66^ Although morphology suggested this direction, the lack of CK7 and HLA-G expression prevented confirmation in our study.^67^ The established CD43-GFP reporter iPSC lines revealed the onset of hematopoiesis within hemanoids on day 6 of cytokine stimulation. This aligns with a report on remodeling EHT ex vivo, describing the appearance of initial CD43+ cells on days 6-8.^35^ CD43+ cells expanded from their area of origin and migrated within hemanoids before being released into the supernatant from day 14 onward. Hematopoiesis was initially organized in blood islands morphologically resembling the YS vascular plexus.^68^ Macrophages were the only HC type in the extravascular connective tissue, which might support the assumption that primitive macrophages, unlike primitive RBCs and megakaryocytes, originate from a monopotent progenitor.^69,70^ In older hemanoids, the endothelial barrier was disrupted, and HCs became evenly distributed between the vessel rudiments and the connective tissue in parallel to their continuous release into the supernatant. We speculate that the growing HC mass within blood islands and the increasing mechanical pressure disrupt the endothelial layer, rather than the transendothelial migration of hematopoiesis. Further investigation is needed, such as using an endothelial reporter alongside the CD43 reporter, to clarify how HCs migrate from blood islands into connective tissue and whether similar mechanisms may be relevant in vivo, such as the transition of YS-derived EMPs to the FL. During continuous culturing, the stromal compartment became highly organized, while hematopoiesis decreased, aligning with a reduced release of HCs into the supernatant. Our observations closely overlap with recent data from the human YS, showing that the hematopoietic-to-stromal cell ratio decreases from young (CS10) to old (CS22) YS. The authors suggested that a loss of stromal support between 6-8 wpc in the YS leads to apoptosis and depletion of remaining hematopoiesis through terminal differentiation.^10^ Despite the contribution of various cell types to the hemanoid, exclusively CD43+ HCs were released into the supernatant. We hypothesize that developing HCs lose their adherent properties and can emigrate from the hemanoid scaffold, while the framework of non-hematopoietic cells remains captured within. CD43 itself has been proposed to mediate anti-adhesive properties of HCs.^35^ Combining the CD43R-iPSC line with approaches to disrupt different receptor-ligand interactions (described below) may shed light on the adhesion capacities of extraembryonic HCs during their maturation.

The presence of monocytes, mast cells, and granulocytes, along with expression of *MYB*^49,71,72^, and fetal globin genes, indicates the presence of a more advanced hematopoietic wave that at least corresponds to EMP-derived hematopoiesis.^3,4^ Protein analysis confirmed about 70% HbF and 20% embHb in hemanoid-derived cRBCs.^22^ Further induction of AGM-derived hematopoiesis appears to be limited. Adult β-globin and increased expression of negative regulators of embHb and HbF (SOX6 and BCL11A) were not detectable.^4,43–45^ Lymphoid potential was restricted to a few CD3+ cells, and we could not confirm expression of recently described signature genes for AGM-derived HSCs^9^ within a single hematopoietic cluster. Overall, our results suggest that predominantly an intermediate EMP program is present in day 16 and day 28 hemanoids, rather than a definitive AGM-derived program capable of producing cells resembling in vivo-generated HSCs. The development of AGM-derived hematopoiesis is significantly influenced by WNT signaling^50^, NOTCH signaling^51,52^, and HOX pathways.^9,12^ The experimental conditions used in our study do not specifically target these pathways. We suggest that hematopoiesis in the hemanoid system becomes exhausted after 5 weeks because i) AGM-derived hematopoiesis is not induced, ii) a complete FL environment is absent, and iii) support from the microenvironment diminishes. It would be very interesting to see if AGM-derived hematopoiesis could be induced by modifying HOX pathways, as recently described.^73^

Our study aimed to identify niche factors that influence hematopoietic cell fate. ST data revealed expression of distinct adhesion molecules on HC populations. Besides inter-hematopoietic cell contacts, we identified hepatoblasts and stromal cells as potential interaction partners. During human embryogenesis, the FL acts as the second major site of hematopoiesis, with crosstalk between hematopoietic and FL cells.^74^ In vivo, liver rudiments develop as a diverticulum from the floor of the embryonic gut around 21 dpc (CS10).^5^ Unexpectedly, we consistently identified clusters of hepatoblasts near blood islands. Based on their transcriptional profile, they might support hematopoiesis by providing growth factors (IGF-2, EPO) and their involvement in iron, lipid, and vitamin B12 metabolism. We recently demonstrated the importance of lipids for RBC differentiation.^33^ EPO expression by endodermal cells of the human YS or FL has been reported,^10,24,75^ as has IGF-2 production by FL cells, and its importance for HSC expansion.^76,77^ Interestingly, hepatoblasts contribute to the production of ECM (vitronectin, fibronectin, fibrinogen, and the laminin-ß3 chain). While laminin mediates cell attachment, migration, and tissue organization during embryogenesis^78–80^, the expression of vitronectin and fibronectin in FL is discussed to contribute to FL colonization by YS EMPs and HSCs.^81^ Moreover, studies on primary human YS hematopoiesis suggest that endodermal vitronectin and YS fibronectin interact specifically with distinct integrins on HCs^10,56^ and thereby modify HSC function, expand the HSC pool, and contribute to long-term HSC quiescence.^10,82,83^ We confirmed gene expression for all corresponding integrin subunits in the EryMK cluster, which might indicate similar mechanisms in hemanoid-derived hematopoiesis.

In the hemanoid system, 3D-organization and stromal cell formation are essential for HC production.^22^ The CD43R-iPSC line revealed strong attachment of HCs to the stromal compartment, consisting of MSCs, fibroblasts, and various ECM molecules. Based on ST data, and in line with published data on YS hematopoiesis^10^, attachment of HCs to the ECM might involve the collagen receptor CD36, and the fibronectin receptors α4β1- and αvβ1-integrin. The supportive role of MSCs or ECM in erythroid differentiation is well established, as they are used to enhance hematopoietic or erythroid growth in culture.^82,84^ Several top DEGs in the Stromal cluster are known to regulate cellular development and differentiation. Interestingly, we observed upregulated TGF-ß pathways across different cell types, including stromal cells (e.g. *TGFB2, BMP5*^58^, *FMOD*^59^), myeloid cells (Fibrillin, modulating TGF-ß bioviability^60,61^), and ERyMK progenitors (*TGFB1*). TGF-ß2 regulates cell growth, migration, and differentiation during embryogenesis. As TGF-ß signaling also regulates a wide range of biological processes in HSCs,^85^ TGF-ß pathways might influence hematopoiesis and erythropoiesis inside hemanoids. TGF-β1 production by megakaryocytes, with supportive effects on terminal RBC differentiation, has been reported.^86,87^

## Future Perspectives

In this study, we explored the enormous potential of iPSCs as a source for in vitro modeling of human hematopoietic and erythroid development. We demonstrated that hemanoids reflect human YS-derived extraembryonic erythropoiesis, spanning the undifferentiated iPSC to the enucleated RBC stage, and provided insight into tissue organization that might affect RBC development. Since extraembryonic hematopoiesis of human origin is not available for repeated experiments due to ethical concerns and physical inaccessibility of embryonic material, reproducible culture systems provide an important tool for studying the earliest physiological stages of hematopoiesis. Unlike other established systems, the hemanoid system not only produces erythroid precursors but also generates enucleated RBCs in sufficient quantities for further functional studies. Therefore, it fills a gap by offering insight into the structure and function of the earliest embryonic RBCs. This includes their membrane composition, blood group antigen expression, biomechanical properties, and oxygen-binding capacities. Such analyses are crucial for evaluating the potential of iPSC-derived cRBCs for clinical applications in transfusion medicine. While cRBCs closely resemble native cells, observed minor differences may arise from their different developmental origins rather than cultural conditions. The characteristics of iPSC-derived RBCs might be advantageous for transfusions in preterm infants, as the oxygen-binding capacities of embryonic and fetal hemoglobin may protect them from cell-toxic effects caused by high oxygen levels and free radical injury.^88^ Because preterm infants often require only small RBC volumes (less than 10 mL), producing sufficient amounts of cRBCs becomes feasible. The successful expansion of 3D-hemanoids in a bioreactor has recently been demonstrated.^28,29,89^ Utilizing autologous iPSCs can further advance personalized medicine approaches, including CRISPR/Cas9-based genome editing to model or treat hematological disorders. Consequently, the hemanoid system may serve as a platform for future clinical translation to study RBC diseases or for de novo RBC production. Resources provided by our study, including the CD43R-iPSC line, refined protocols for ST analysis of small spheroids, ultrastructural STEM images, and ST data, will support future research in the vital field of developmental hematopoiesis.

## Limitations of the study

Our study is limited by the resolution of the Visum 10X ST system. This constrains more precise identification of cell clusters and the detection of rare events such as the emergence of HSCs. To obtain single-cell-level information and more accurately define cell populations and their characteristics, a scRNA-seq analysis of the hemanoids is planned for a future study. The cell interaction mechanisms identified in this study need to be confirmed in further research and examined for their functional importance.

## Resource availability

### LEAD contact

Requests for further information and resources should be directed to and will be fulfilled by the lead contact, Isabel Dorn.

### Materials availability

The CD43-GFP reporter iPSC line generated in this study is available upon request from the lead contact with a completed materials transfer agreement. Usage must comply with the ethics approval on which the patient’s consent was based and exclude commercial use.

### Data and code availability

The ST data have been deposited in NCBI GEO as GSE324601 and are publicly available as of the date of publication. STEM images have been deposited in Dryad as [Dataset DOI: 10.5061/dryad.69p8cz9hz] and are publicly available as of the date of publication.

This paper analyzes existing, publicly available scRNA-seq data, accessible at E-MTAB-11673, GEO: GSE144024, GEO: GSE144024, and E-MTAB-7407.

This paper does not report original code.

Any additional information required to reanalyze the data reported in this paper is available from the lead contact upon request.

## Acknowledgments

We thank M. Sundl for assistance with immunohistochemistry and sample preparation for ST analysis; K. Hingerl for assistance with STEM analysis; the Core Facility Bioimaging (M. Absenger) for live-cell imaging support; the Core Facility Molecular Biology (B. Gallé and N. Schweintzger) for ST sample processing; S. Trajanoski for his input on ST data analysis; and E. van den Akker for providing Sani-003A-iPSCs. This research was funded by the Austrian Science Fund (FWF), Grant-DOI 10.55776/I6572 to I.D. and 10.55776/PAT9611123 to G.M. Figures were created with BioRender.

## Author contributions

I.D. and A.A. designed the study; M.A. and A.A. performed iPSC culturing and ex vivo erythropoiesis; A.A. generated the GFP reporter iPSC line; M.A., D.B., G.H., and J.F. performed immunohistochemistry; D.K., M.A., and I.D. performed STEM analysis; A.A., M.A., G.M., and I.D. designed and performed Spatial Transcriptomics analyses; A.A. performed the bioinformatics analysis; A.R., P.S., J.F. and I.D. monitored and supervised the study; C.B. and P.S. performed project administration and provided resources; I.D. and A.A. analyzed and interpreted the experiments, and wrote the original manuscript; all authors reviewed and edited the final version of the manuscript.

## Methods

### Human material and cell lines

The study was approved by the local ethics committee at the Medical University of Graz in line with the Declaration of Helsinki (27-165ex14/15). Three human iPSC lines derived from erythroblasts were used (UBTi001-A (PEB-AL#6)^30^; CM1^32^, Sani-003A^31^) and cultured at 37°C and 5% CO_2_ in Stem MACS iPS Brew XF medium (#130-104-368, Miltenyi Biotech) on 6-well tissue culture plates coated with Matrigel® (#354277, Corning). Cells were mechanically split every 6-7 days and supplemented with 10 µM Rock Inhibitor (#130-106-538, Miltenyi Biotech). Human K-562 cells (ACC-10, DSMZ) were cultured in RPMI-1640 medium (#11875093, Gibco) supplemented with 10% fetal bovine serum (FBS) (#S0615, Biochrom) and 1% Penicillin Streptomycin (PS) (#15070063, Gibco). HEK293T cells (#300189, CLS) were maintained in DMEM high-glucose (#D5671, Sigma-Aldrich) with 10% FBS, 1% PS, and 25 mM HEPES (#15630056, Gibco).

### Hemanoid formation and erythroid differentiation of iPSCs

Hematopoietic and erythroid differentiation of iPSCs was performed as recently described ^22,23^ and illustrated in Figure 1. For EB generation, iPSC colonies were detached from the tissue culture well using 1mg/mL collagenase type IV (#17104019, Gibco). Cell aggregates were seeded on ultra-low-binding plates (#15277905, Nunclon Sphera, Thermo Scientific) and cultivated for 5 days in hESC medium without bFGF.^90^ Thereafter, spherical EBs were transferred onto six-well tissue culture plates in STEMdiff™ APEL™ 2 medium (#5270, StemCell Technologies), supplemented with 5% Protein-Free Hybridoma Medium (#12040-077, ThermoFisher Scientific), 100 ng/mL SCF (#300-07, Peprotech), 5 ng/mL IL-3 (#200-03, Peprotech), and 3 U/mL EPO (Erypo, Janssen Biologics B.V.). The medium was changed weekly. Within a few days, EBs adhered to the tissue culture plate and formed self-organized 3D structures, termed hemanoids. The size and the morphology of the hemanoids were assessed by microscopy (EVOS M5000, ThermoFisher Scientific). Hemanoids were characterized at different maturation stages by flow cytometry and immunohistochemistry. For flow cytometry characterization, hemanoids were digested into a single-cell suspension using 0.4 IU/mL Collagenase B (#11088815, Roche) (2 h at 37°C, 5% CO2) and Cell dissociation buffer (#13151014, Gibco) (10 min at RT) as described.^91^ Single cells released from hemanoids into the supernatant were repeatedly harvested to determine cell counts, characterize them by flow cytometry, and assess hematopoietic colony formation in semisolid media.

For erythroid differentiation, released single cells were repeatedly harvested from the supernatant and cultured for 18 days in an established erythroid differentiation protocol.^33^ Culture medium consisted of Iscove’s Modified Dulbecco’s Medium (IMDM) (#FG0465, Biochrom) containing 5% Octaplas^®^ LG (Octapharma), 10 µg/mL insulin (#91077C, Sigma-Aldrich), 330 µg/mL transferrin (#T101-5, BBI Solutions), 1% PS, and from day 8 onwards 4mg/dL choles-terol-rich lipids (#L4646, Sigma-Aldrich). Cells were stimulated as follows: Day 0 to day 8: 100 ng/mL SCF, 5 ng/mL IL-3, 3 U/mL EPO, and 10^-6^ M hydrocortisone (OHC) (#H2270, Sigma-Aldrich); Day 8 to day 11: 100ng/mL SCF and 3 U/mL EPO; day 11 to day 21: 3 U /mL EPO. Cell numbers and cell vitality were counted in a Malassez counting chamber after staining with trypan blue (#T8154, Sigma-Aldrich). Hematopoietic and erythroid differentiation were monitored by flow cytometry and microscopic evaluation of cytospin preparations after staining with May-Gruenwald-Giemsa (#102103, Hemafix, Biomed) and neutral benzidine (#D9143, o-Dianisidine, Sigma-Aldrich) for the detection of hemoglobin. At least 300 cells were enumerated under the microscope (Axioscope, Zeiss). In some experiments, day 18 cells were filtered through an Acrodisc WBC syringe filter (#AP-4851, Pall Corporation) to obtain the pure enucleated portion of cultured RBCs.

### Flow cytometry

Flow cytometry analysis was performed on a CytoFLEX^®^ flow cytometer (Beckman Coulter) using the CytExpert software 2.4. The following antibodies were used to stain the cells throughout hematopoietic and erythroid differentiation: CD34-PE (#A07776, Beckman Coulter), CD43-APC (#560198, BD Biosciences), CD45-PC7 (#IM3548, Beckman Coulter,), CD45-FITC (#IM3454808, BD Biosciences), CD36-FITC (#B49201, Beckman Coulter), CD235a-FITC (#B49206, Beckman Coulter), CD49d-APC (#B01682, Beckman Coulter), CD71-PE (#555537, BD Biosciences), and CD233 (Band3)-PE (#9439PE, IBGRL). Dead cells were excluded by 4’,6-Diamidino-2-phenylindol (DAPI) (#D3571, ThermoFisher Scientific) staining. Graphs were partially generated using FlowJo (TreeStar, v10).

### Colony formation

Single cells released from the hemanoid into the supernatant were collected and plated in triplicate (2,500 cells/dish) on MethoCult (#H84434, StemCell Technologies) coated 35mm dishes (#27100, StemCell Technologies). After 10-14 days, the dishes were scored using light microscopy (Primovert; Zeiss). Colonies were classified into burst-forming unit-erythroid (BFU-E), colony-forming unit-erythroid (CFU-E), colony-forming unit-granulocyte/erythrocyte/monocyte/megakaryocyte (CFU-GEMM), colony-forming unit-macrophage (CFU-M), and colony-forming unit-granulocyte/macrophage (CFU-GM).

### Immunohistochemistry

Hemanoids were washed 2x with Dulbecco’s Phosphate Buffered Saline (DPBS) (#14190-094, Gibco™), fixed for 1 hour in 4% paraformaldehyde (#1.04005.1000, Merck Millipore), and embedded in paraffin with Excelsior™ (Thermo Fisher). Paraffin-embedded hemanoids were sectioned (10µm sections) using a Thermo ScientificTM rotation microtome HM355s (Fisher Scientific). Before staining, antigen retrieval was performed in a microwave for 2 x 20-minute cycles in citrate buffer (pH 6). UltraVison™ Quanto Detection System HRP (#TL-015-QHD, Epredia™) was used for antibody detection using the Horseradish peroxidase (HRP) system. Briefly, after being washed 3x with PBS, the slides were incubated for 10 minutes with UltraVision™ Hydrogen Peroxidase Block, washed 3x again with PBS, and incubated for 5 minutes with UltraVision™ Protein. Primary antibodies, rabbit anti-human CD43 (#MA5-16339, ThermoFisher Scientific), recombinant rabbit anti-human CD144 (VE-Cadherin) (#MAB-44374, ThermoFisher Scientific), rabbit anti-human HBE1 (#PA5-106357, ThermoFisher Scientific), rabbit anti-human Laminin beta-1 (#PA5-27271, Thermo Fisher), mouse anti-human vimentin (#MAB3400, Merck, Millipore), rabbit anti-human AFP (#145501-AP, Proteintech), mouse anti-human HBG1 (#66168-1-Ig, Proteintech), mouse anti-human CD68 (#14-0688-82, Thermo Fisher Scientific), mouse anti-human CD163 (#DB 045, DB Biotech), mouse anti-human CD14 (#60253, PTGlab, Proteintech), mouse anti-human 235a (#M0819, Dako, rabbit anti-human EPX (#ab238506, Abcam), rabbit anti-human myeloperoxidase (#GA511, Dako), mouse anti-human mast cell tryptase (#IR640, Dako), mouse anti-human Integrin beta 3 (#ab9509, Abcam), mouse anti-human CD20 (#GA604, Dako), rabbit anti-human CD3 (#GA503, Dako), mouse anti-human CK7 (#MS-1352-P, ThermoFisher Scientific), mouse anti-human HLA-G (#557577; BD Biosciences) were diluted in antibody diluent (#TA-125-ADQ, Epredia™) according to manufacturer’s instructions and the slides were incubated for 60 minutes. Thereafter, the slides were incubated for 10 minutes with Primary Antibody Amplifier Quanto, washed again 3x, and incubated for 10 minutes in the dark with HRP Polymer Quanto (light sensitive). Coloring was performed for 5 minutes using a mixture of DAB Quanto Chromogen and Substrate (one drop of chromogen in 1 ml substrate) (brown color) or for 10 minutes using 4 drops of AEC Substrate (red color) (#ab64252, Abcam). Counterstaining to identify the tissue morphology was performed using modified hematoxylin (H&E, #8947.1, Roth) for 2 minutes. The whole staining process was performed in a humidified chamber at RT. IHC slides were scanned using a digital slide scanner (Slideview VS200, Olympus, Tokyo, Japan) equipped with an LED source (Excelitas Technologies, X-Cite Xylis, Mississauga, Canada) and a CMOS camera (2304=×=2304, ORCA-Fusion C14440-20UP, 16-bit, Hamamatsu, Japan). Analysis software for scanned slides was Olympus OlyVIA 3.4.1.

### Scanning Transmission Electron Microscopy (STEM)

Hemanoids were fixed in in 0.1M cacodylate buffer (2.14% w/v Dimethylarsinic acid sodium salt trihydrate (#820670, Sigmal-Aldrich) in Aqua bidest, pH 7.4) supplemented with 2.5% glutaraldehyde (#16200, Electron Microscopy Sciences) and 2% paraformaldehyde (#1.04005.1000, Merck Millipore) for 3h, and then post-fixed in 2% osmium tetroxide (#19110, Electron Microscopy Sciences) for 2h at RT. After dehydration in a graded series of ethanol (50%-100%), tissues were infiltrated in propylene oxide (#149620010, ThermoFisher Scientific) for 1h, followed by stepwise infiltration in TAAB Embedding Resin (TER) (#T004, TAAB Laboratories Equipment Ltd., UK): 50% v/v TER in propylene oxide for 3h at RT, followed by 66% v/v TER (overnight, 4°C), and finally pure TER (3 h, 45°C). Embedded tissues were transferred to embedding molds (#10590, PELCO) and polymerized for 48h at 60°C. Semithin sections (1µm) were cut with glass knives (#7890-04, Leica Microsystems) and stained with Toluidin blue solution (1% w/v Dinatriumtetraborate (Sigma-Aldrich, #106306) and 1% w/v Toluidine blue (#R1727, Agar Scientific) in Aqua Bidest). Slides were assessed using a BX41 light microscope (Olympus). Ultrathin sections (70 nm) were cut with a UC 7 Ultramicrotome (Leica Microsystems, Austria) and a diamond knife (#2302, Diamond Knife DiATOME Sciences Services), placed on pioloform-covered grids (#R1275 Agar Scientific) (#G200H-Cu and G2010Cu, Sciences Services), and stained with 1% platinum blue (EMS, USA, #22407) for 15 min and 3% lead citrate (#16707235, Leica Microsystems) for 5 min. Electron micrographs were taken using a Tecnai G2 transmission electron microscope (Thermo Fisher Scientific, Netherlands) with a Gatan Ultrascan 1000 charge-coupled device (CCD) camera (-20°C; acquisition software: Digital Micrograph, Ametek Gatan, Germany; and Serial EM). The acceleration voltage was 120 kV. To image large areas of hemanoids at high resolution, scanning transmission electron microscopy (STEM) imaging mode on a field-emission scanning electron microscope (ZEISS FE-SEM Sigma 500) with an acceleration voltage of 15 kV, in combination with ATLAS TM (version 5.2.2.15, ZEISS), was used.

### Plasmid-AAV design and cloning

The AAV vector plasmid was cloned into the pAAV-MCS plasmid (#240071, Agilent Technologies) containing inverted terminal repeats from AAV serotype 2 (AAV2), with a maximal packing capacity of 4,7 kb. The donor plasmid was assembled by standard Gibson assembly (**Table S1)** of the NotI HF (#R3189S, NEB) linearized plasmid backbone using NEBuilder® HiFi DNA Assembly Mastermix (#E2621L, NEB). The constructed plasmid contains the right (RHA) and left homology arms (LHA), each 300 bp, homologous to the DNA flanking the spCas9 cut site (***AAVS1_sgRNA: GGGGCCACUAGGGACAGGAU***), 5’ splice acceptor to ensure the exact splicing after transcription, T2A a self-cleaving peptide, puromycin-resistant cassette with bovine growth hormone (bGH) polyadenylation signal, followed by a long (2179 bp) CD43 promotor region which drives the expression of the fused green fluorescent protein (GFP) and the SV40 polyadenylation signal.

### AAV production

Human embryonic kidney 293T cells (HEK293T cells) were expanded to a total of at least 130 million cells in DMEM high glucose (Sigma-Aldrich) supplemented with 4 mM L-glutamine (#G7513, Sigma-Aldrich), 1mM Sodium pyruvate (#11360070, ThermoFisher), 10% FBS (Biochrom), 1% PS (Gibco), 25 mM HEPES (Gibco), and 1 mM Sodium butyrate (#B5887, Sigma-Aldrich). 13 million cells were seeded per individual 15 cm dish one day before transfection. At about 70–80% confluency, cells were transfected using 5 µg/mL polyethyleneimine (#23966, Polysciences). For transfection of a total of 10 plates, 60 μg AAV donor plasmid (*pAAV6_AAVS1_CD43_GFP*) and 220 μg helper plasmid pDGM6^92^ (was a gift from David Russell, #110660, Addgene) were mixed with PEI in Opti-MEM (#3798570, Gibco). The mixture was incubated at RT for 15 minutes, then added dropwise to the media and carefully swirled. Cells were incubated in a humidified 37°C incubator for 72 hours. AAV6 viral particles were harvested and purified using the AAVpro Purification Kit (#6666, Takara) following the manufacturer’s instructions. Viral particles were stored in aliquots at - 80°C till further use. The copy number/µl was determined by ddPCR (QX200, Biorad).

### Electroporation of iPSCs

After digestion with Accutase (#T8154, Sigma-Aldrich), the iPSC single-cell suspension was electroporated using the Lonza 4D Nucleofector (program CA-137) and the P3 Primary Cell Nucleofection Kit (#V4XP-3024, Lonza). We have electroporated as few as 300,000 cells per condition using the electroporation strips holding 20 µl. The RNP complex (Cas9 + sgRNA mixed in a 1:2.5 molar ratio, Cas9 Nuclease V3, #1081059, Gibco) was prepared at 25°C for 15 minutes before electroporation and scaled down based on the number of electroporated cells. After electroporation, the cells were transduced with 5,000 – 10,000 AAV vector genomes/cell and incubated at 37°C and 5% CO_2_. Cells were seeded as single cells into six-well plates containing pre-warmed antibiotic-free XF media (Miltenyi). After 48h of incubation, small colonies were scored under the microscope. Puromycin selection was initiated after cells reached about 40-50% confluency. Puromycin (#A11138-03, Gibco) was used at concentrations ranging from 0.1 µg/mL to 0.5 µg/mL, depending on the iPSC line. Puromycin-resistant cells were picked and clonally expanded before further approval of successful gene targeting.

### Characterization of iPSCs edited clones

Genomic DNA was purified from the clones using QuickExtract™ DNA Extraction Solution (#QE09050, Epicentre). Briefly, the mixture of cells and extract solution 25 µl was vortexed for 15 seconds, incubated at 65°C for 6 minutes, vortexed for 15 seconds, and incubated again at 98°C for 2 minutes. The DNA was amplified using an in-out PCR (one primer within the introduced DNA sequence, the other primer outside of the homology arms) (**Table S2**). The amplicon was Sanger-sequenced (Eurofins Genomics) to confirm the knock-in at the AAVS1 harbor locus.

### Live cell imaging and immunofluorescence microscopy

The emergence of hematopoietic CD43-GFP+ cells within adherent hemaloids was observed by live-cell imaging using a Nikon HCS Ti2 Eclipse (Celesta V2) microscope. Hemanoids were analyzed at different time points. Image acquisition settings were optimized for GFP detection. The time frame was set to one image every 10, 30, or 60 minutes for 24 or 72 hours. Samples were imaged with a 20x objective (NA 0.75), and excitation was provided by a 488 nm laser. Emission was collected using MXR10018 1^st^ 4000 KG Phometrics BSI Express CMOS camera/PMT. Images were acquired using NIS Elements C software with Jobs (v5.42.07). Additional observation of CD43-GFP+ cells was done with an EVOS M5000 microscope (ThermoFisher Scientific).

### Spatial transcriptomics sample processing and sequencing

10 to 18 hemanoids generated from the PEB-AL#6 iPSC line were pooled on days 16 and 28, respectively (step II). The hemanoids were washed 2x with PBS (Gibco), fixed for 60 minutes using 4% paraformaldehyde, and embedded in paraffin using Excelsior™ (ThermoFisher Scientific). For spatial transcriptomics, the hemanoids after RNA quality assessment (DV200) were cut into 5 µm thick sections using a rotation microtome HM355s (ThermoFisher), and mounted within the capture areas of the Visum Spatial Gene Expres-sion Slide (#PN-1000189, 10X Genomics). The slide was dried at 40°C on a heat plate. Tissue deparaffinization, H&E staining, imaging, and decrosslinking were performed according to the 10X Visium Spatial Gene Expression for FFPE guideline (10X Genomics, Deparaffinization, H&E Staining, Imaging & Decrosslinking, CG000409 Rev C). H&E images were taken using a Leica Aperio ScanScope^®^ AT imaging system at x40 magnification and the Aperio ImageScope software (v12.4.6.5003). Probe extension and library construction steps using the Visium FFPE Reagent kit (PN-1000361) and the Visum Human Transcriptome Probe kit (#PN1000363) followed the 10X user guide Visium Spatial Gene Expression Reagent Kits for FFPE (CG000407 | Rev D). Tissue slides had an average DV200 of 28%. The coverage area was estimated with the Loupe Browser v7 (10X Genomics, Pleasanton, California, U.S.). Sequencing was performed with the recommended read mode: read 1: 28 cycles; i7 index read: 10 cycles; i5 index read: 10 cycles; and read 2: 50 cycles on an Illumina NextSeq2000.

### Processing of spatial RNA sequencing reads

After sequencing, the reads were aligned to the human genome (hg38) using the Space Ranger pipeline (v4.0.1, 10X Genomics) with default parameters. Space Ranger was also used to align paired histology images with the positions of mRNA capture spots on the Visium slides. The raw UMI count matrix, images, spot-image coordinates, and scale factors were imported to the Seurat R package (v5.1.0)^93^ for downstream data processing. In brief, we first performed quality control (QC) to remove low-quality spots based on metrics including total UMI counts, the number of detected genes, and the percentage of mitochondrial gene expression. Spots with unusually low or high gene counts, low UMI counts, or high mitochondrial content were excluded from further analysis. Following QC, we used SCTransform to normalize and scale the data and identify variable genes. Dimensionality reduction was performed using RunPCA, and the first 20 principal components were used for downstream analyses. We applied FindNeighbors to these components, followed by FindClusters to cluster the ST spots at a resolution of 1.0. Finally, we used RunUMAP on the same 20 principal components to visualize the data in two dimensions. Differentially expressed genes (DEGs) for each cluster were identified using the FindAllMarkers or FindMarkers functions in Seurat with default parameters, comparing gene expression within each cluster to all remaining clusters. DEG analysis was performed using a pairwise Wilcoxon Rank-Sum test between spots within each cluster and all other spots in the dataset. The DotPlot function was used to illustrate the expression pattern of selected genes for different cell types or conditions.

Cell type annotation and label transfer were performed using the Azimuth package v0.5.0^94^ with the fetal development reference data^42^, followed by manual annotation using known marker genes for clusters that showed additional diversity in gene signatures.

We used Harmony v1.2.0^55^ to integrate and batch-correct the ST data of FFPE samples (days 16 and 28) with published scRNA-seq datasets. Namely, the scRNA-seq data from yolk sac (E-MTAB-11673^54^, and GEO: GSE144024^53^), and fetal liver (GEO: GSE144024^53^, and E-MTAB-7407^11^). Prior to integration, each dataset was individually preprocessed using the Seurat pipeline, including normalization, identification of highly variable features, and scaling. After quality control and preprocessing, the datasets were merged into a single Seurat object. Following the merge, we repeated the standard analysis steps: we performed principal component analysis (PCA) with RunPCA, constructed a shared nearest-neighbor graph using FindNeighbors, identified clusters using FindClusters, and visualized the integrated data using RunUMAP. Gene ontology (GO) enrichment analysis for the selected cell clusters was performed in R, using the enrichGO function from the clusterProfiler v4.6.2^95^ package and the “org.Hs.eg.db” database. Gene set enrichment analysis (GSEA) was performed on gene lists identified by the FindMarkers function as statistically significant. The gseGO function from the clusterProfiler package v4.6.2^95^ with default parameters was used. The selected pathways were visualized using the R package ggplot2 v4.0.1.^96^ For comparison, we have also utilized the fgsea v1.32.4^97^ package in R.

### Statistics

Data are presented as mean ± standard deviation (SD) unless otherwise stated. Raw data were tested for normality of distribution, and statistical analyses were performed using a two-tailed unpaired t-test, a two-tailed Mann–Whitney test, a Wilcoxon rank sum test, a two-way ANOVA for the line graphs, and a Kruskal–Wallis test with multiple comparison tests, depending on the dataset. GraphPad Prism 10.4.1 (GraphPad Software, San Diego, CA, USA) or R (R-Studio, R 4.2.0) was used for statistical analyses.

## Supplemental information

The Supplemental information (Document S1) includes Figures S1 - S13, Tables S1 and S2, and supplemental references.

