## Supplemental Figures and Tables for "Self-organized hemanoids derived from human iPSCs create a niche that produces definitive extraembryonic hematopoiesis"

### Supplemental information

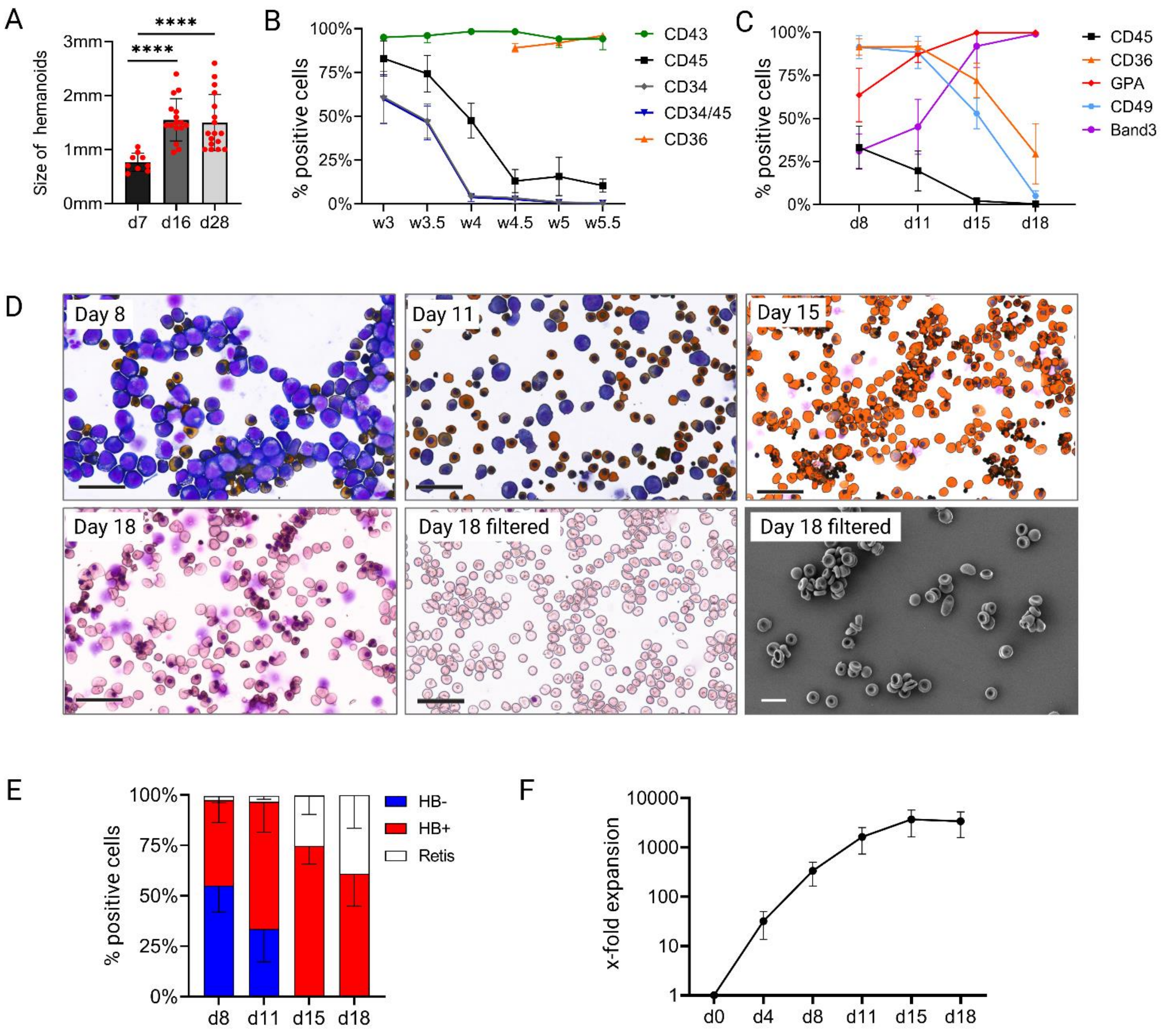


**Figure S1. Characterization of single cells released from hemanoids into the supernatant and their further erythroid differentiation, related to Figure 1.**

(**A**) Bar graph showing the size of hemanoids (diameter of complexes without stromal layer) determined by microscopy in millimeters (mm) on days 7 (n=9), 16 (n=16), and 28 (n=18); (mean ± SD, ****p<0.001, unpaired t-test). (**B**) Time course of cell surface marker expression on cells released from the hemanoids into the supernatant, analyzed by flow cytometry (week 3 to week 5.5 of phase I), (n=3; mean ± SD). Due to the limited number of cells, CD36 measurements started in week 4.5. (**C**) Flow cytometry analysis of cell surface markers expressed from days 8 to 18 during the erythroid differentiation (phase II), (n=6; mean ± SD). (**D**) Representative images of cytospin preparations costained with May-Grünwald Giemsa (MGG) and neutral benzidine to detect hemoglobin (scale bar 50 µm). Day 18 cells were stained only with MGG for a more valid evaluation of enucleation. Lower middle figure: Filtered enucleated cells from day 18. Lower right figure: Representative scanning electron microscopy image of filtered iPSC-derived enucleated cultured RBCs, displaying the typical morphology of RBCs (scale bar: 10 µm). (**E**) Differential counts of stained cytospin samples from days 8 to 18 of erythroid differentiation (hemoglobin-negative erythroid precursors (HB-), hemoglobin-positive erythroid precursors (HB+), enucleated reticulocytes (Retis) (n=6; mean ± SD). (**F**) Cumulative expansion of erythroid cells from days 0 to 18 of phase II (n=6; mean ± SD). Data in B, C, E, and F were derived from 3 different iPSC lines as biological replicates.

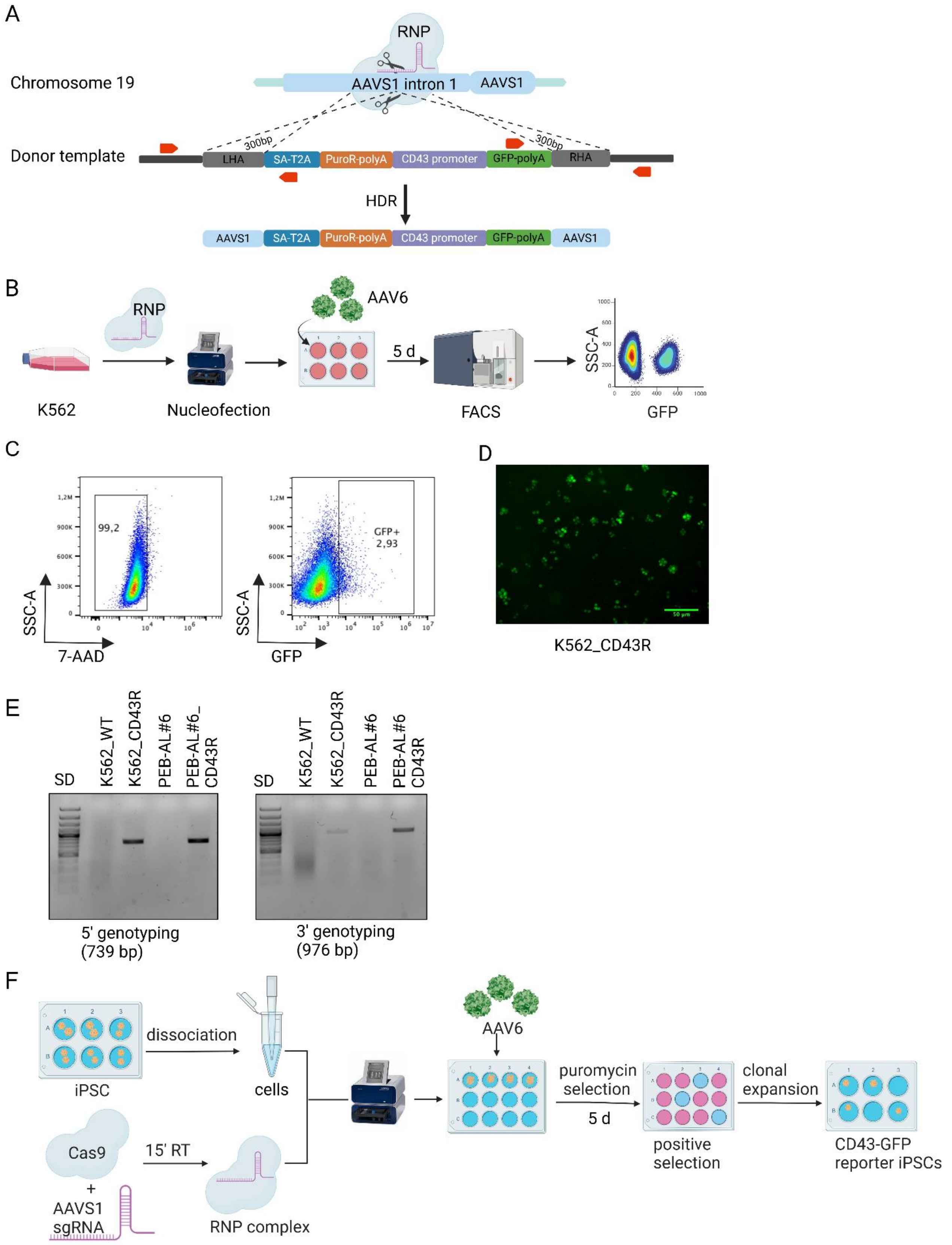


**Figure S2. CRISPR/Cas9-mediated CD43-GFP reporter iPSC line generation, related to Figure 2.**
(**A**) Schematic presentation of the HDR-based knock-in strategy into the AAVS1 harbor locus on chromosome 19 and of the donor template (left to right): left homology arm (LHA), splice acceptor sequence (SA) and self-cleaving peptide sequence (T2A), puromycin-resistant cassette with a polyadenylation tail, CD43 promoter region tagged with a DNA sequence encoding GFP and another polyadenylation tail, and right homology arm (RHA). Red arrows indicate the primer binding sites for in-out PCR genotyping of the 5' and 3' HDR junctions. (**B**) Workflow for CRISPR/Cas9-mediated generation of K562-CD43-GFP reporter (K562_CD43R) cells and their enrichment via flow cytometry-activated cell sorting (FACS). K562 cells were electroporated with the RNP

complex (Cas9+sgRNA) and transduced with AAV6 carrying the donor template. After 5 days, GFP+ cells were enriched via fluorescent-activated cell sorting (FACS). (**C**) Flow cytometry pseudocolor plots of K562 cells 5 days post-AAV6 transfection. Live cells were gated by excluding 7-AAD-positive cells (left), and GFP expression was then evaluated, indicating efficient transfection (right). (**D**) Fluorescence microscopy image showing CD43-GFP expression in the K562 cell line after the transfected cells were enriched via fluorescent-activated cell sorting (scale bar: 50 µm). (**E**) Gel images picturing the 5' (739 bp) and 3' (976 bp) ends after PCR genotyping. The correct integration of the donor DNA template into the AAVS1 locus was confirmed by PCR amplification using construct-specific primers. Shown are the correct integration of the CD43 promoter constructs into the AAVS1 locus of K562_CD43R cells, and PEB-AL#6_CD43R iPSCs, in comparison to unmanipulated cells. (**F**) Workflow for generating the CD43-GFP reporter iPSC lines using the CRISPR/Cas9 RNP method and AAV6 as a template carrier to knock in the desired DNA sequence into the AAVS1 locus.

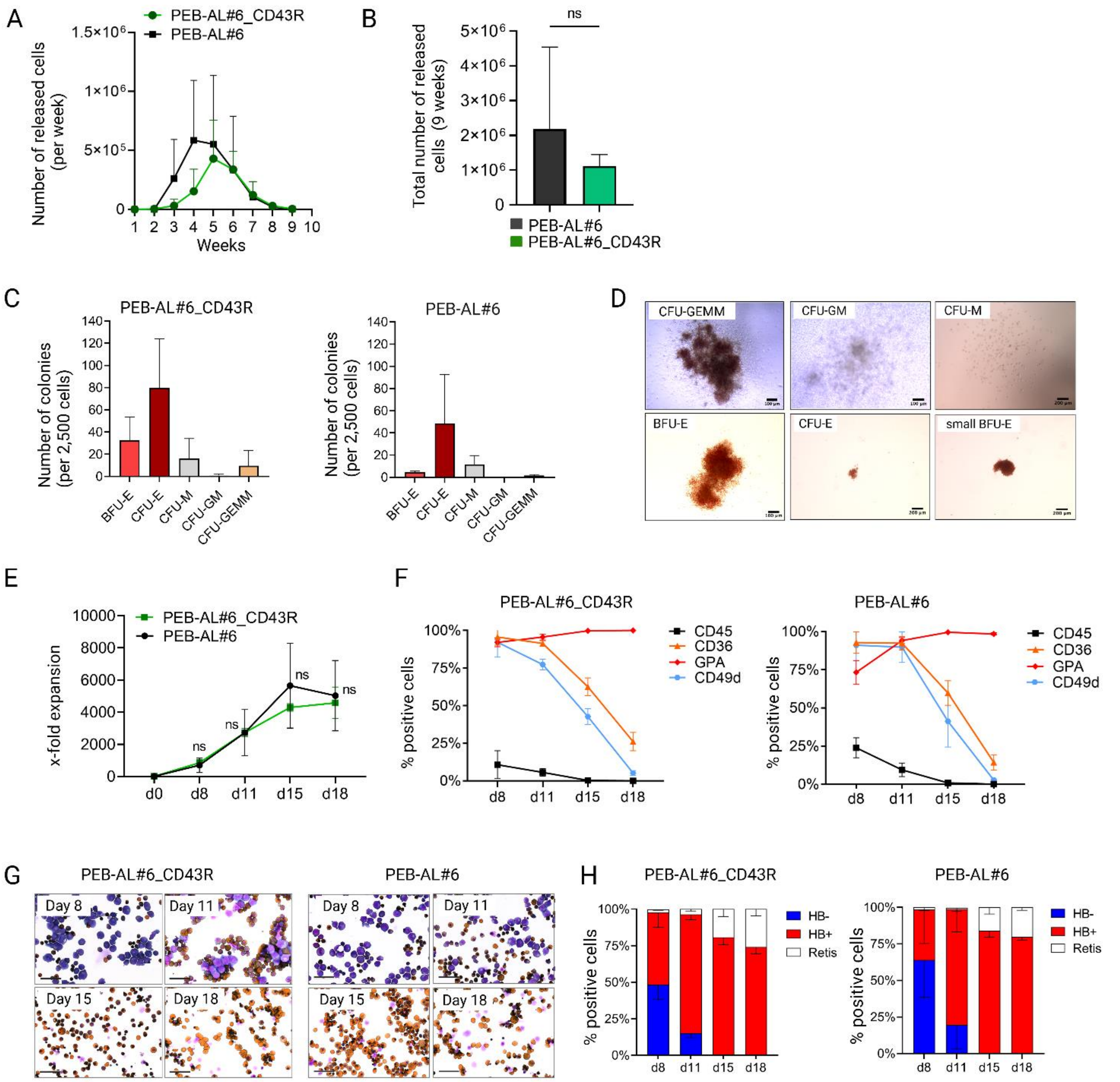


**Figure S3. The hematopoietic and erythroid potential of the PEB-AL#6_CD43R iPSC line compared to the unmodified PEB-AL#6 iPSC line, related to Figure 2.**
(**A**) Hemanoids from PEB-AL#6 and PEB-AL#6_CD43R iPSCs were maintained in culture for nine weeks (phase I). Shown is the number of single cells released from the hemanoids into the supernatant per week (weeks 1 to 9; n = 3; mean ± SD). (**B**) Total number of single cells released from PEB-AL#6 and PEB-AL#6_CD43R hemanoids into the supernatant over the nine-week period (n=3; mean ± SD, unpaired t-test, ns not significant). (**C**) The released single cells (between weeks 3 and 3.5) were subjected to a colony-forming assay in semisolid media. Hematopoietic colonies were counted and scored after 14 days (n=3; mean ± SD). (**D**) Representative microscopic images showing the morphology of the colonies as scored: Colony-forming unit–granulocyte, erythrocyte, monocyte, megakaryocyte (CFU-GEMM), Colony-forming unit-granulocyte, macrophage (CFU-GM), Colony-forming unit-macrophage (CFU-M). Burst-forming unit–erythroid (BFU-E), and colony-forming unit–erythroid (CFU-E). CFU-Es showed signs of a more primitive nature (pale red color, small size) (scale bars: 100 µm and 200 µm). (**E**) Cumulative expansion of PEB-AL#6 and PEB-AL#6_CD43R derived cells during further erythroid differentiation (phase II) (n=3; mean ± SD). (**F**) Flow cytometry analysis of cell surface markers expressed from days 8 to 18 during the erythroid differentiation of PEB-AL#6 and PEB-AL#6_CD43R cells (n=3; mean ± SD). (**G**) Representative images of cytospin preparations prepared during erythroid differentiation (phase II), costained with May-Gruenwald Giemsa and neutral benzidine (scale bar 50µm). (**H**) Differential counts of stained cytospin samples from days 8 to 18 of erythroid differentiation (hemoglobin-negative erythroid precursors (HB-), hemoglobin-positive erythroid precursors (HB+), enucleated reticulocytes (Reti) (n=3; mean ± SD).

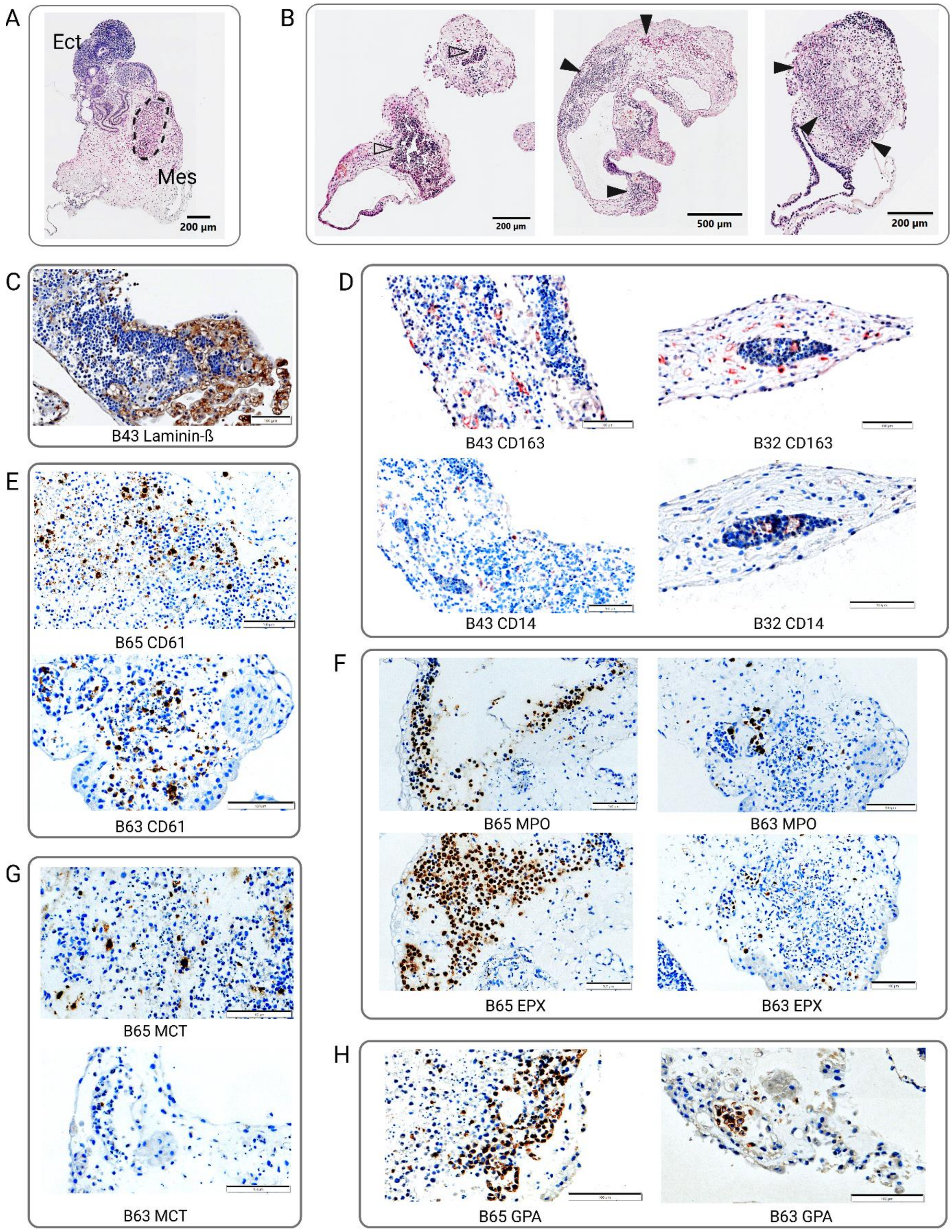


**Figure S4. Confirmation of different hematopoietic cell types inside hemanoids by HE- and specific antibody staining, related to Figure 3.** (**A**) Representative brightfield image of a hemanoid showing ectodermal (Ect) structures comparable to neural crest tubes and a mesoderm-derived (Mes) stromal compartment with embedded blood island (dashed black line) (scale bar: 200 µm). (**B**) Hemanoid sections showing hematopoiesis in vessel-like structures (left, white arrows) or inside the stromal compartment (middle & right, black arrows) (scale bars 200 µm and 500 µm). (**C**) Hemanoid B43, co-stained by horseradish peroxidase (HRP) for Laminin-ß. (**D**) Tissue sections from hemanoids B43 and B32, co-stained for CD14+ monocytes and CD163+ macrophages. Macrophages were also positive for CD68 (not shown). (**E-H**) Tissue sections from hemanoid B63 (18 days) and B65 (4 weeks) co-stained by HRP for cell-type specific antigen expression: (E) CD61+ megakaryocytes and platelets; (F) Myeloperoxidase (MPO) positive granulocytes and Eosinophil Peroxidase (EPX) positive eosinophilic granulocytes; (G) Mast Cell Tryptase (MCT) positive Mast cells; (H) Glycophorin A (GPA) positive erythroid cells. (scale bar in C-H: 100µm)

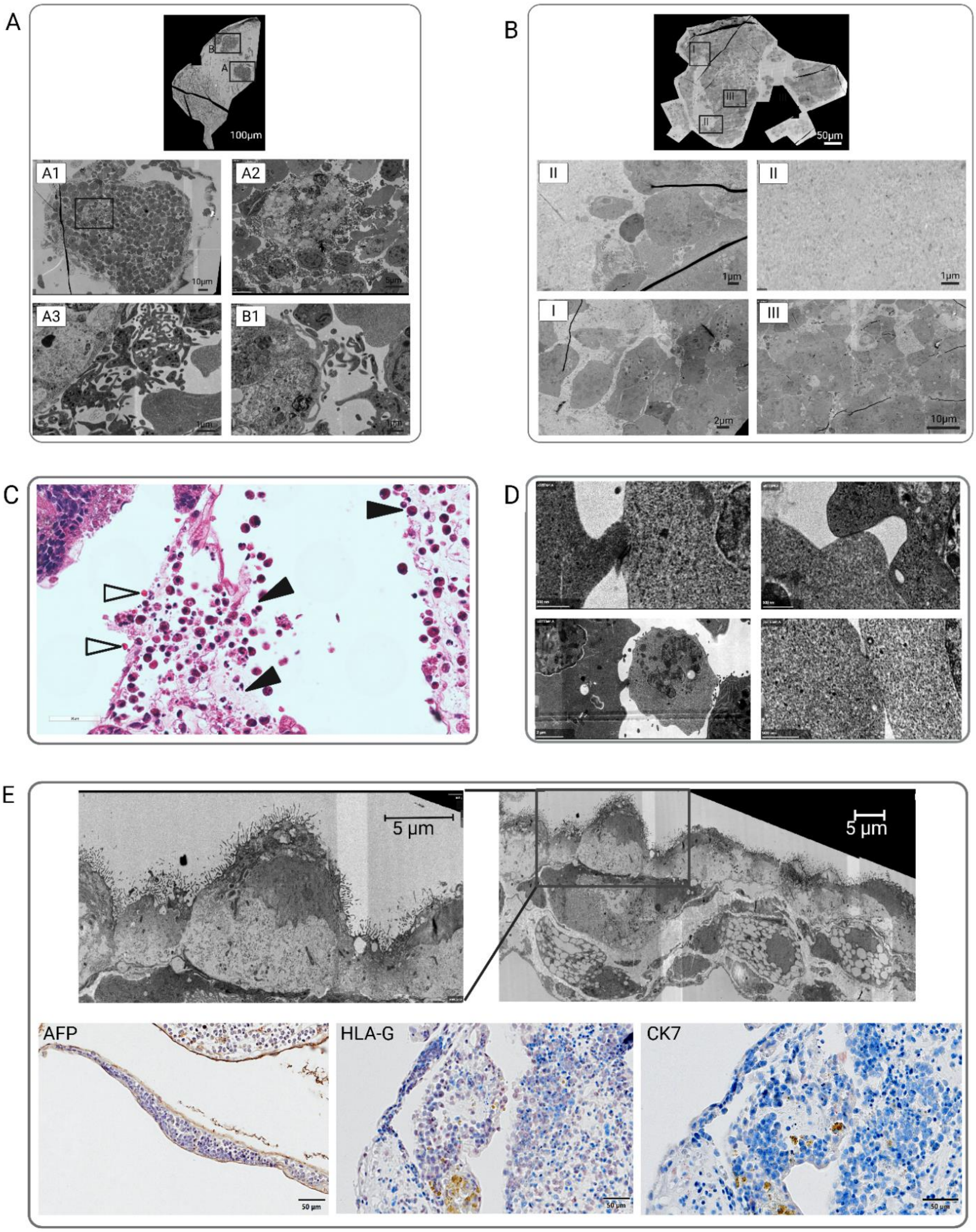


**Figure S5. Characterization of hemanoids by STEM and IHC analysis, related to Figure 4**. (**A**) STEM analyses of a day 17 hemanoid. Shown are increasing magnifications from the identical hematopoietic areas A (A1-A3) and B (B1). Magnifications focus on giant cells (suspected megakaryocytes) generating multiple cytoplasm-derived plasma fragments (platelets). (**B**) STEM analyses of a day 28 hemanoid, with magnifications from areas I-III. Compared to the day 17 hemanoid in S5A and Figure 4, the tissue is more homogeneous, with fewer vessels and HCs detectable. Cavities are filled with collagen fibers. (**C**) HE-staining of day 28 hemanoid sections. Black arrows indicate a damaged endothelial barrier, with HCs entering the surrounding tissue. White arrows indicate mature RBCs (scale bars: 50 µm). (**D**) Cell contacts observed by STEM between erythroid precursors and other cells (scale bars: 0.5 and 2 µm). (**E**) Epithelial layer surrounding hemanoids. Top: STEM images showing the structure of epithelial cells with multiple cell extensions reaching the extracellular lumen. Bottom: HE-stained sections of the epithelial layer, costained for Alpha Fetoprotein (AFP), Human Leukocyte Antigen-G (HLA-G), and Cytokeratin 7 (CK7) (Scale bars: 50 µm). The original day 17 and day 28 STEM image files are accessible at DOI: 10.5061/dryad.69p8cz9hz for an independent view.

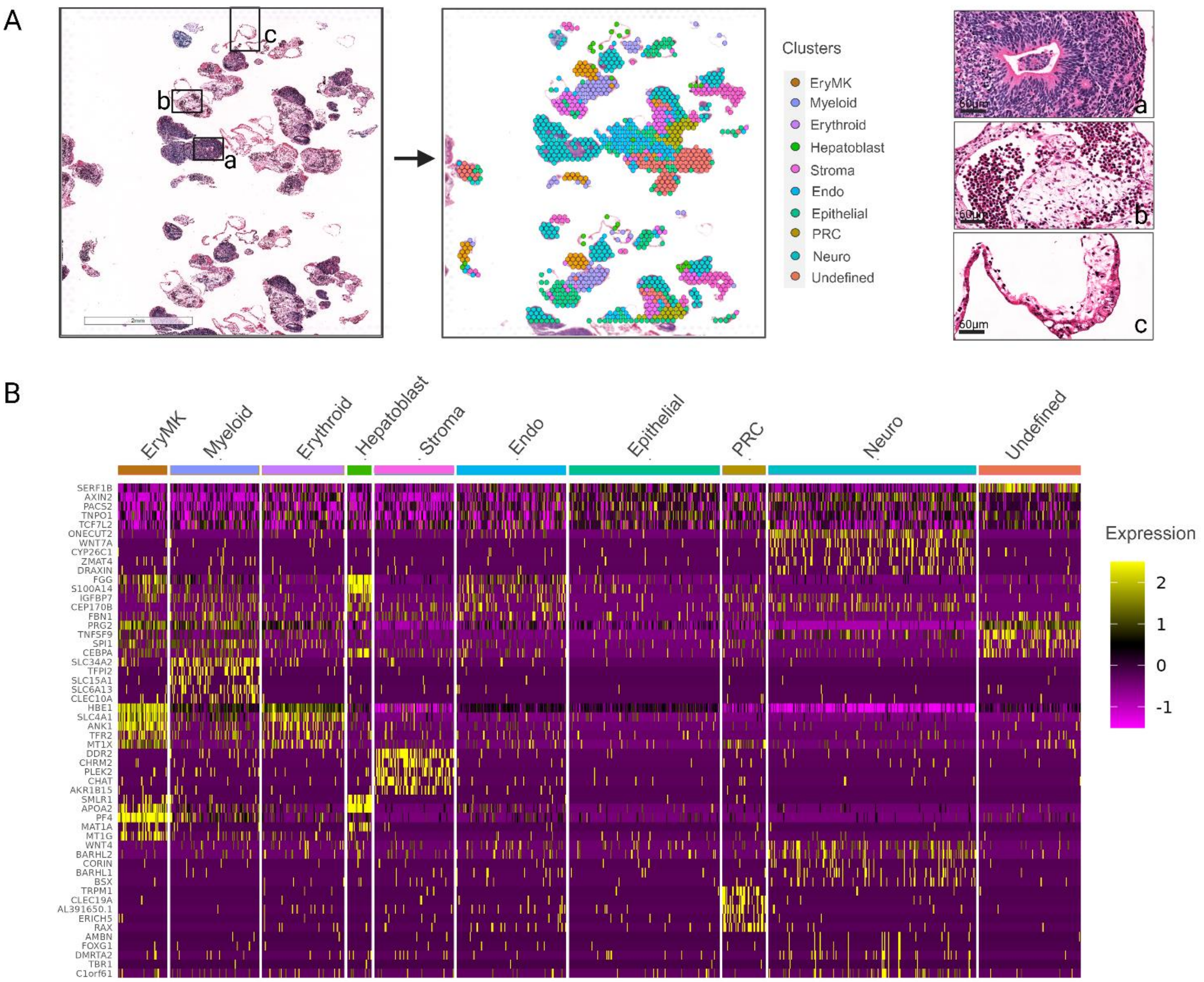


**Figure S6. ST analysis of day 16 PEB-AL#6 hemanoids, related to Figure 5.** (**A**) Left: Microscopic image of the day 16 hemanoid tissue sections (HE-stained, scale bar: 2 mm) subjected to ST analysis before further RNA processing. Middle: Loupe browser projection and spatial clustering distribution. Right: Magnifications from areas a-c in A, with a) Neuro progenitors, b) Stroma including blood islands, c) endodermal-derived epithelial-like cells (scale bars: 60 µm). (**B**) Heatmap showing the average expression level of the top 5 differentially expressed genes for each cluster.

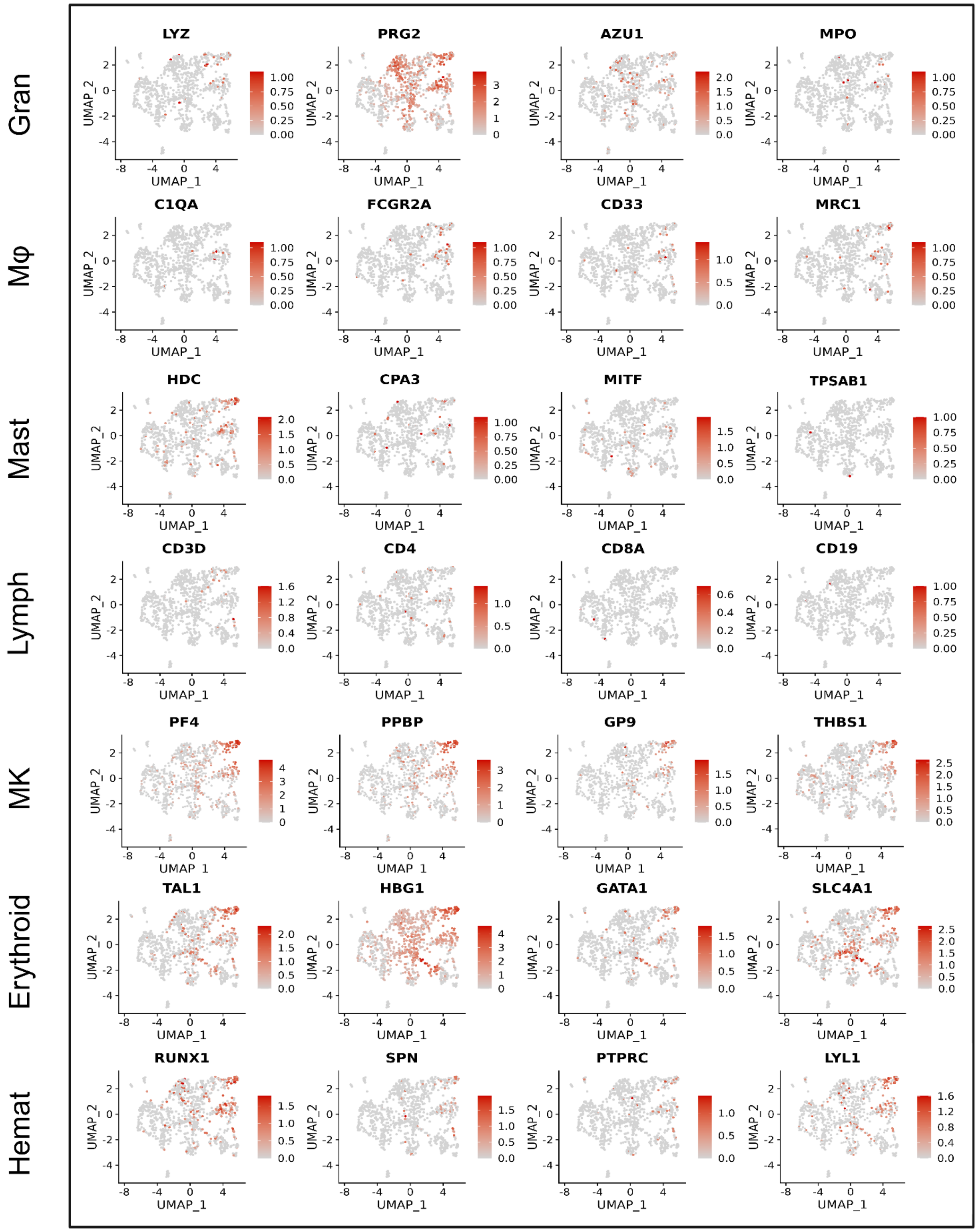


**Figure S7. Expression of marker genes in ST clusters obtained from day 16 hemanoids, related to Figure 5:** Feature plots showing the expression of four representative marker genes for each hematopoietic cell type projected onto UMAP, where the color intensity indicates expression levels (Gran - granulocytes; Mφ - macrophages; Mast - mast cells; Lymph - lymphocytes; MK - megakaryocytes; Erythroid - erythroid cells; Hemat - hematopoietic cells).

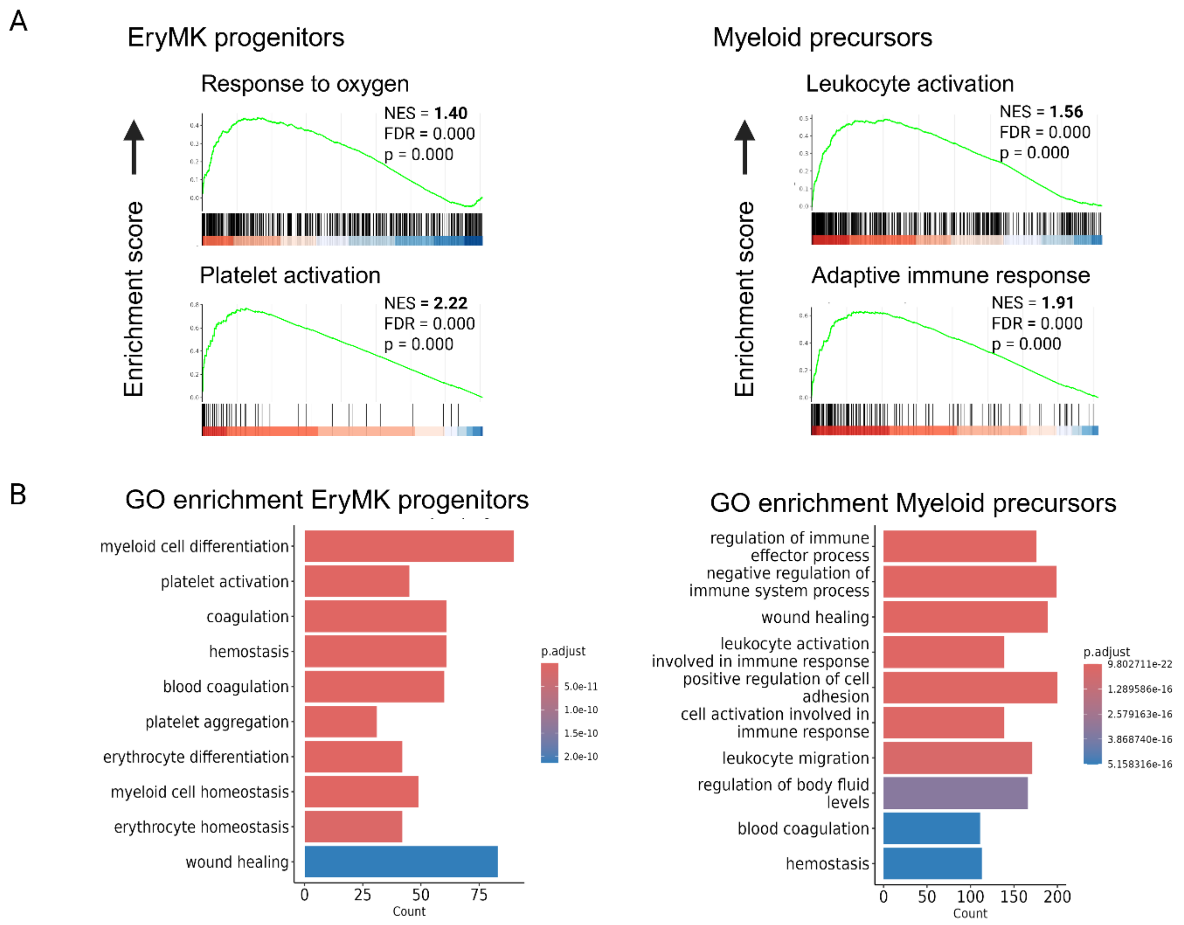


**Figure S8. Enriched pathways in the EryMK and the Myeloid cluster (ST analysis of day 16 hemanoids), related to Figure 5.** (**A**) GSEA plots indicating the enriched pathways in the EryMK cluster (left) and the Myeloid cluster (right). (**B**) Gene Ontology (GO) enrichment analysis was performed on genes identified for the EryMK and Myeloid clusters using the FindMarkers function.

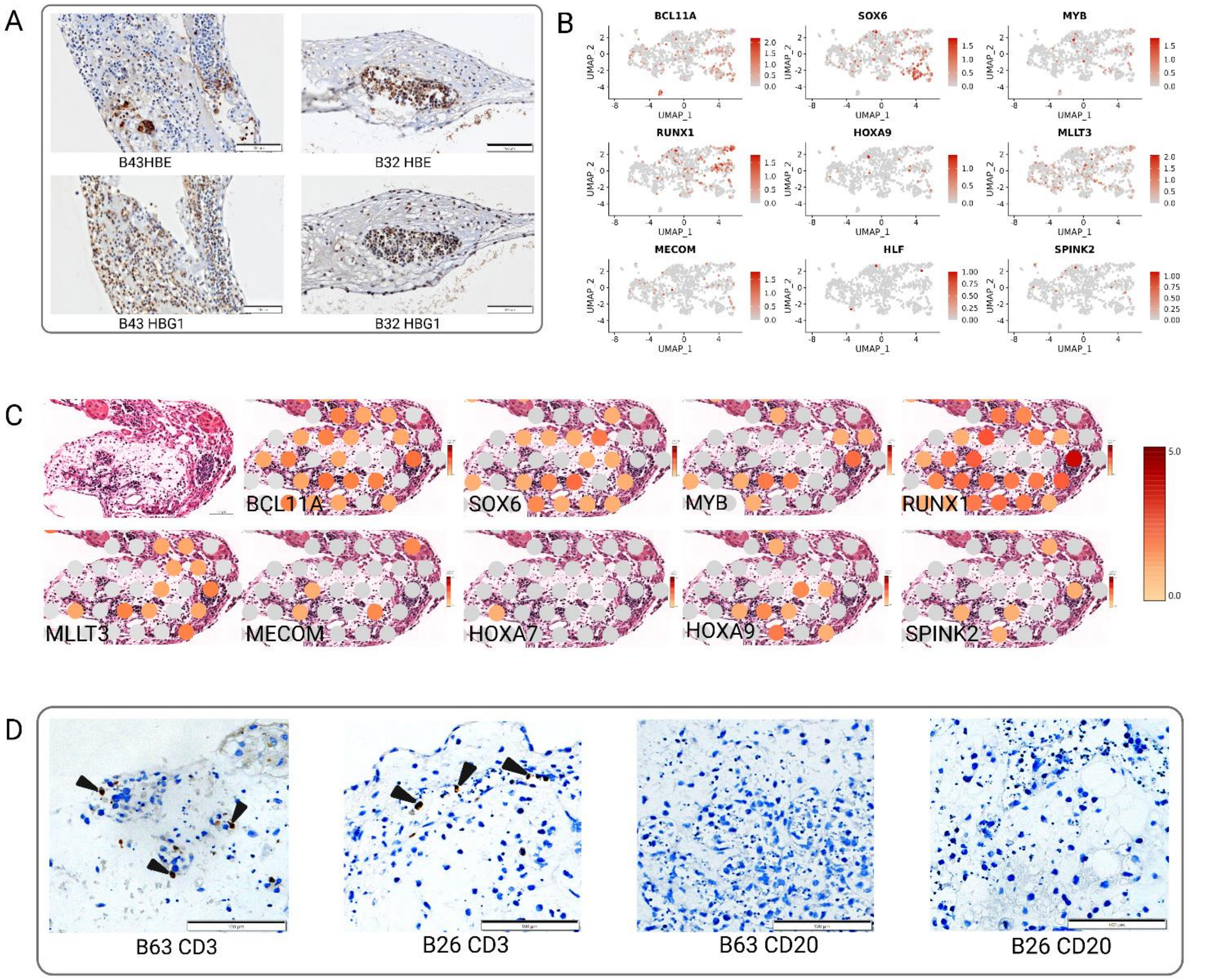


**Figure S9. Expression of developmental wave-specific markers in day 16 hemanoids, related to Figure 6:** (**A**) HE-stained tissue sections from hemanoids B43 and B32, co-stained with HRP for embryonic hemoglobin (HBE) and fetal hemoglobin (HBG1) (Scale bar: 100µm). (**B**) Feature plots from ST analysis showing the expression of *BCL11A*, *SOX6*, *MYB,* and AGM-derived HSC signature genes *RUNX1*, *HOXA9*, *MLLT3*, *MECOM*, *HLF*, and *SPINK2* in d16 hemanoids projected on UMAP. Color intensity indicates expression levels. (**C**) Gene expression overlay on Visium spots from a section of day 16 hemanoids (Loupe browser projection, log2 transformed UMII counts, expression increases from grey to dark red; scale bar: 100 µm). (**D**) HE-stained tissue sections from hemanoids B26 (4 weeks) and B63 (18 days), co-stained with HRP for CD3 (black arrows) and CD20 (Scale bar: 100µm).

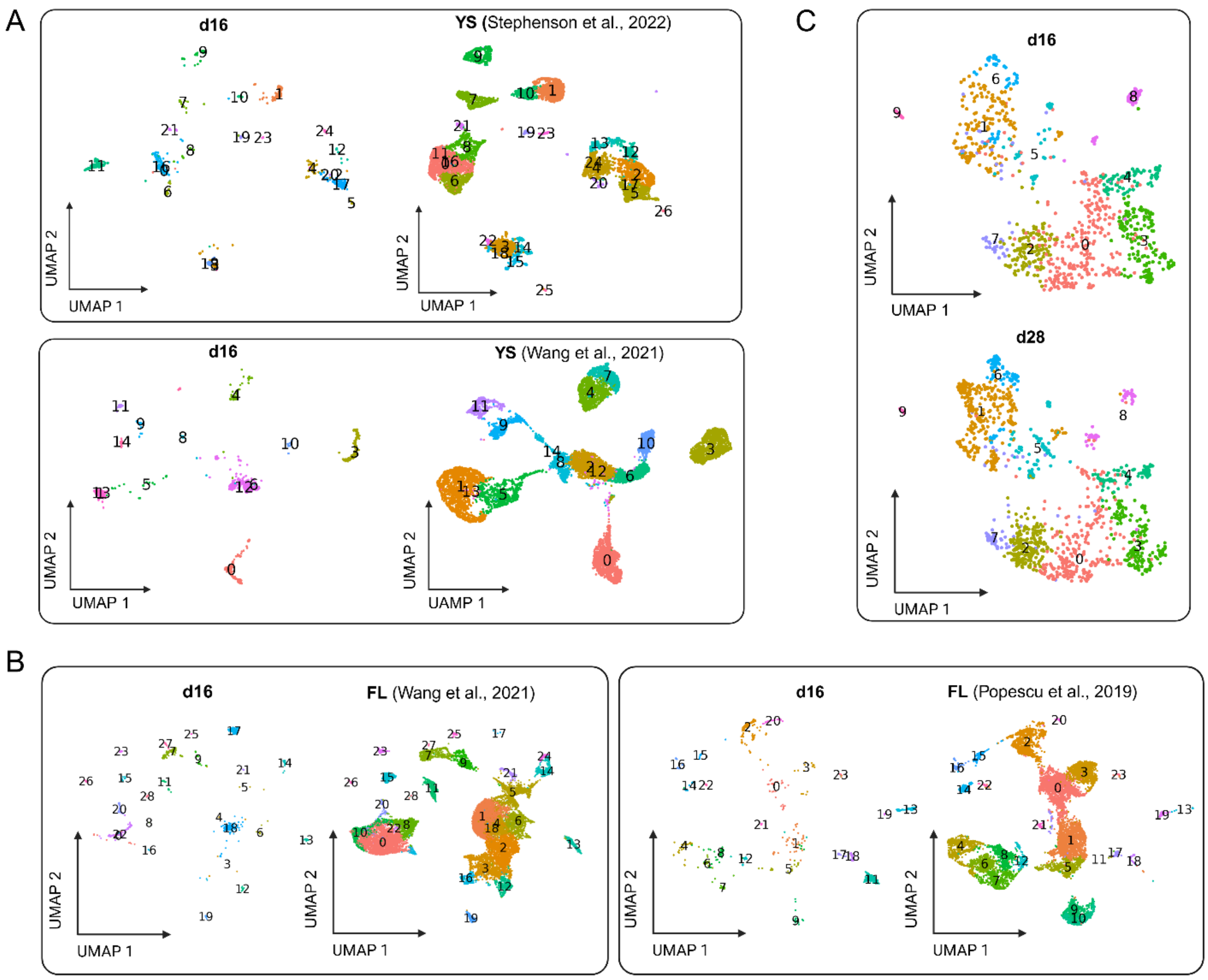


**Figure S10. Comparison of the transcriptional profile (ST data) between day 16 and day 28 hemanoids, and with gene expression data from human YS and FL, related to Figure 6.** **A**) UMAP visualization of Harmony-integrated ST datasets from day 16 hemanoids vs YS-derived scRNAseq data from CS17 (Stephenson et al. [1]) and CS10/11 (Wang et al. [2]). **B**) UMAP visualization of Harmony-integrated ST datasets from day 16 hemanoids vs FL scRNAseq data from CS20/23 (sorted CD34+ cells, Wang et al. [2]), and vs. sorted CD45+ cells from PCWs 8-16 (Popescu et al. [3]). **C**) UMAP visualization of the Harmony integration of day 16 hemanoid vs day 28 hemanoid ST data. Samples in A-C were visualized using the split. by = "sample" parameter in the DimPlot function of Seurat.

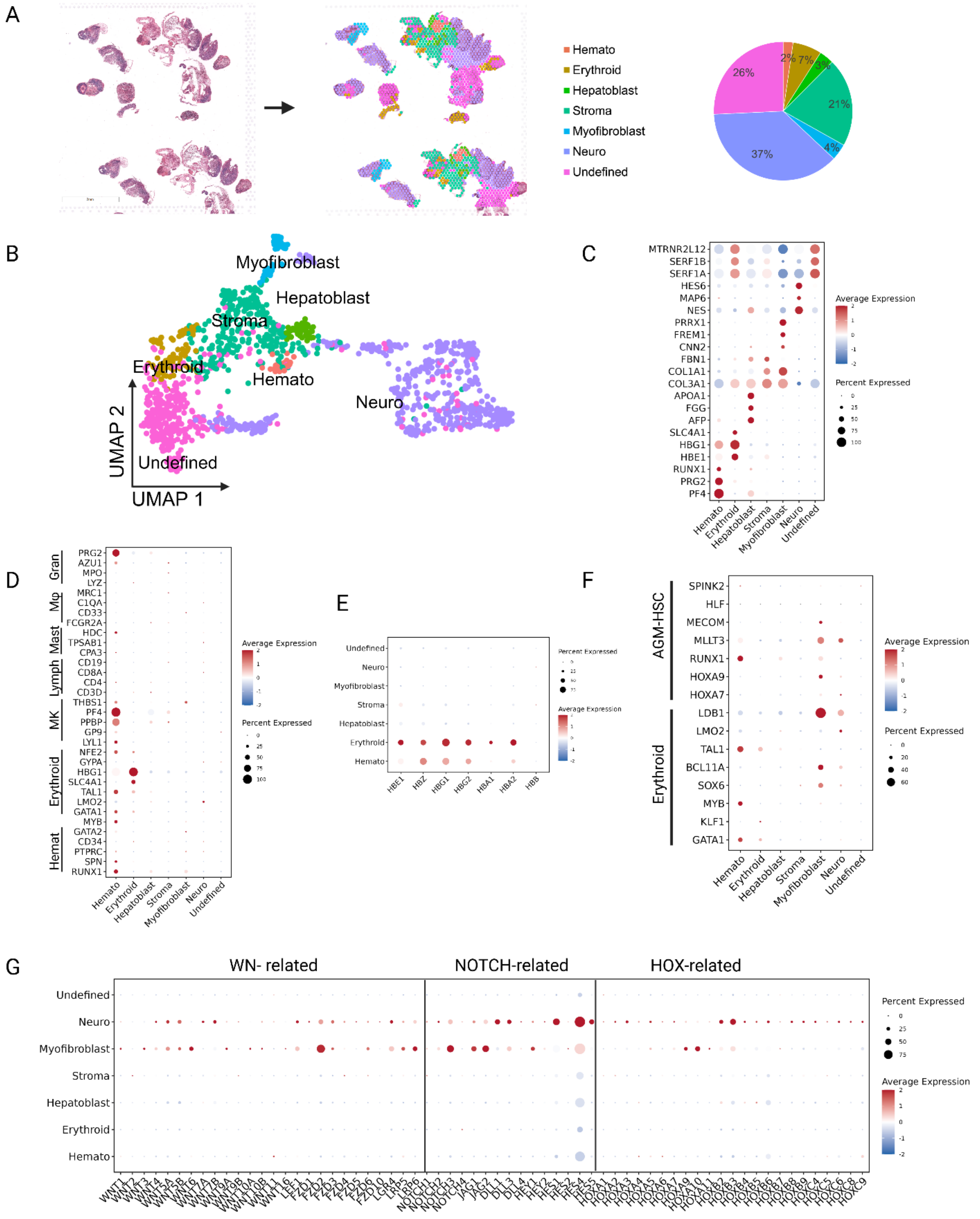


**Figure S11: Spatial transcriptomics (ST) analysis of PEB-AL#6 day 28 hemanoids, related to Figure 6.** (**A**) Microscopic image of the HE-stained day 28 hemanoid tissue sections subjected to ST (left), spatial clustering distribution (middle), and percentage of spots contributing to each of the 7 identified clusters (right). Colors indicate the 7 identified cell clusters based on gene expression. (**B**) UMAP visualization of ST-sequencing data derived from day 28 hemanoid sections. (**C**) Dot plot showing the expression of three representative lineage marker genes in each cluster. Dot size indicates gene expression frequency, and color intensity indicates gene expression level. (**D**) Expression of canonical marker genes for the hematopoietic lineage (Gran (Granulocytes), Mφ (Makrophages), Mast (Mast cells), Lymph (Lymphoid cells), MK (Megakaryocytes), Erythroid (erythroid cells), Hemat (Hematopoietic cells). (**E**) Expression of globin genes encoding the alpha and beta chains of embryonic, fetal, and adult hemoglobin. (**F**) Expression of signature genes for AGM-derived HSCs[9] and of core erythroid transcription factors.[8] (**G**) Expression of genes related to WNT signaling, Notch signaling, and HOX gene clusters.

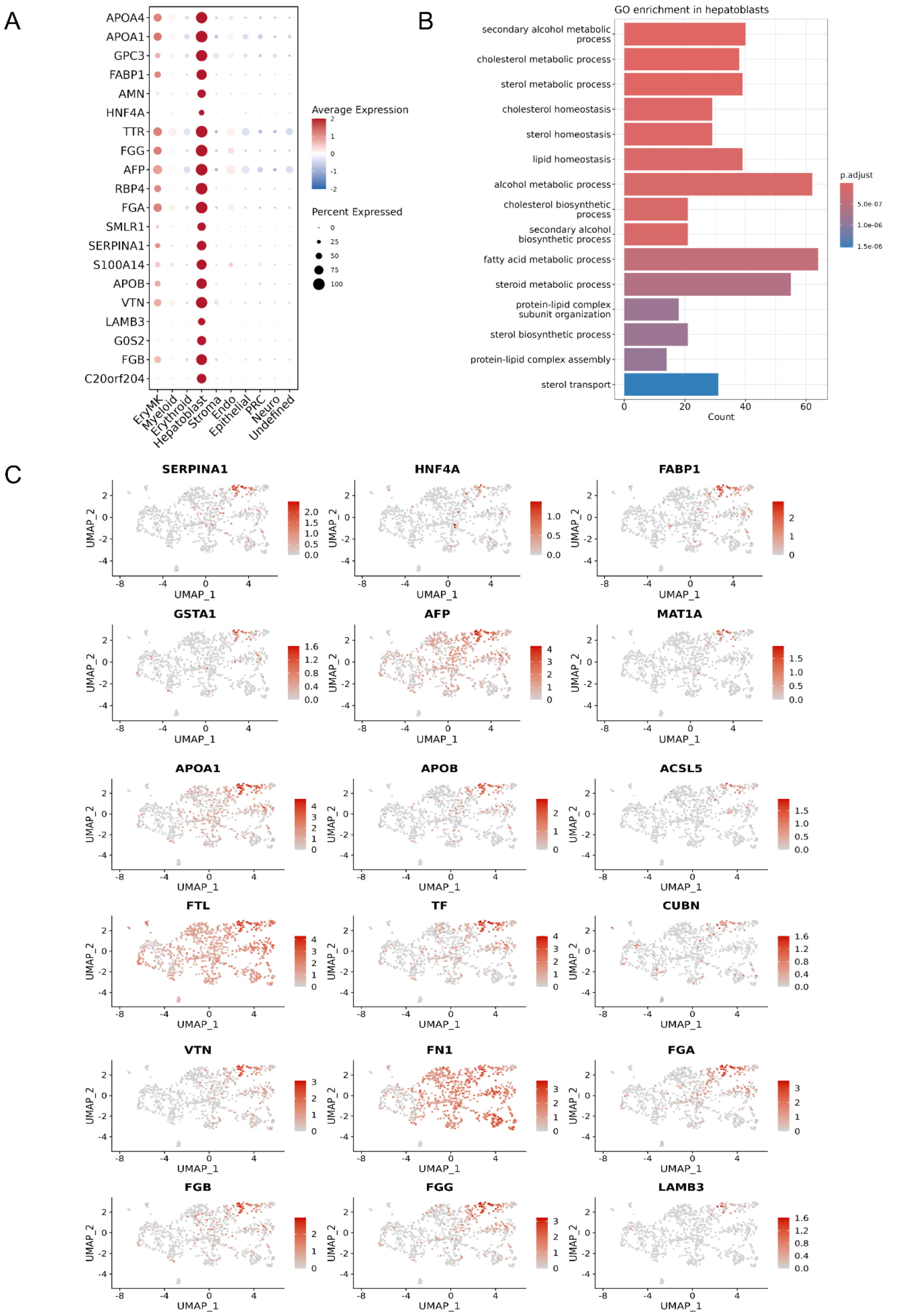


**Figure S12: Gene expression profile of the Hepatoblast cluster obtained by ST analysis of PEB-AL#6 d16 hemanoids, related to Figures 5 and 7. (A)** Dot plot showing the expression of the top 20 variable genes in the Hepatoblast cluster, as calculated by Moran's I. (**B**) Gene Ontology (GO) enrichment analysis of differentially expressed genes in the Hepatoblast cluster. (**C**) Feature plots showing gene expression in the hepatoblast cluster, projected onto UMAP plots, with color intensity indicating expression levels.

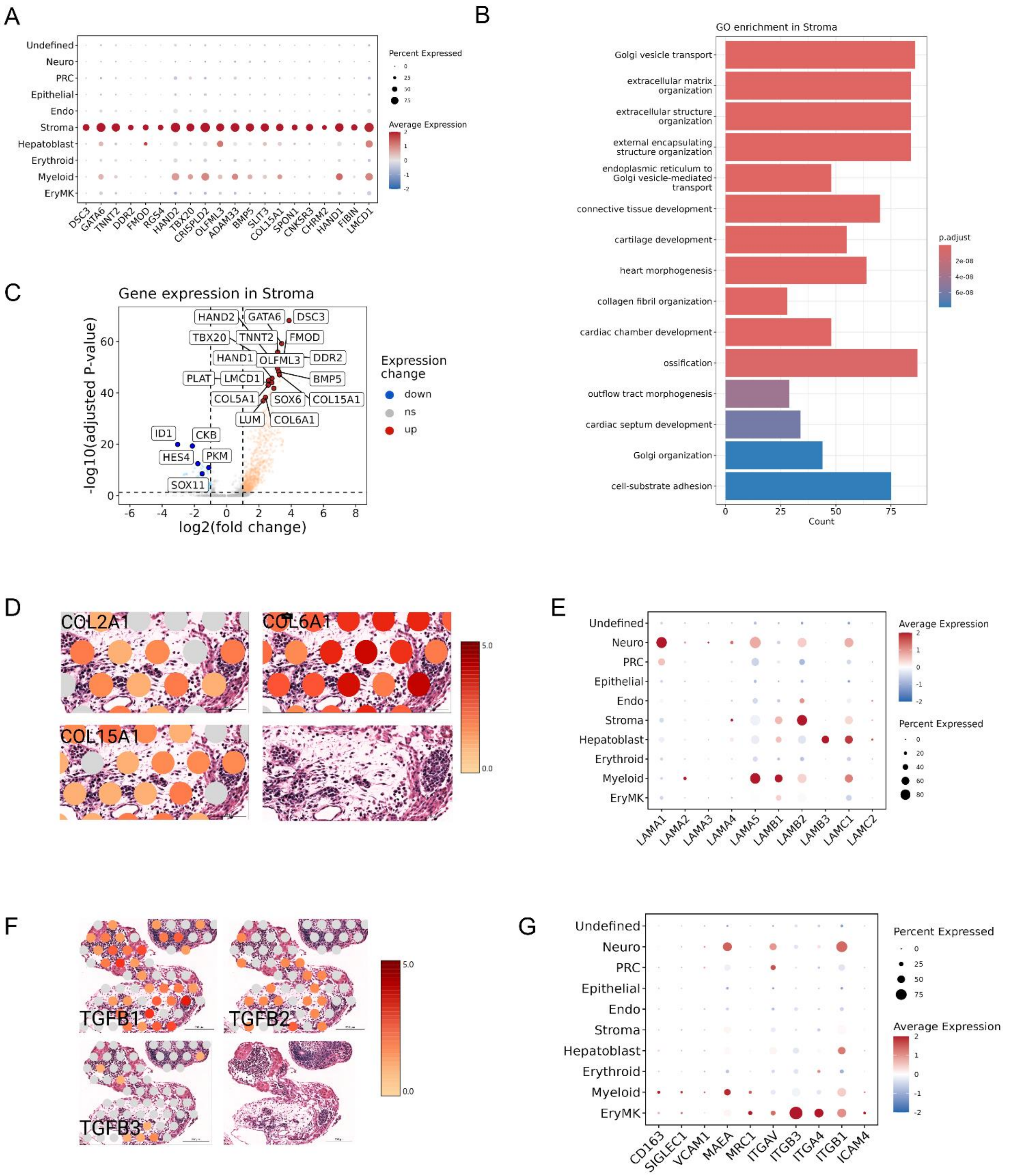


**Figure S13: Gene expression of the Stroma cluster and expression of erythroblastic island markers (ST analysis of PEB-AL#6 day 16 hemanoids), related to Figures 5 and 7. (A)** Dot plot showing the expression of the top 20 variable genes in the Stroma cluster, as calculated by Moran's I. (**B**) Gene Ontology (GO) enrichment analysis of differentially expressed genes in the Stromal cluster. (**C**) Volcano plot showing upregulated and downregulated genes (log2 fold change) in the Stromal cluster. (**D**) Expression overlay of selected collagen genes on Visium spots from a section of day 16 hemanoids. Loupe browser projection (expression increases in $\log_2$ fold from grey to dark red; scale bar: 100 µm). (**E**) Dot plot showing the average expression of Laminin-chain coding genes. (**F**) Expression overlay of *TGFB1-3* genes on Visium spots from a section of day 16 hemanoids. Loupe browser projection (expression increases log2-fold from grey to dark red; scale bar: 200 µm). (**G**) Dot plot showing the average expression of erythroblastic island markers in day 16 hemanoids.

**Table S1. Primers used for Gibson assembly cloning for generating the CD43-GFP-Reporter iPSC lines.** Primers were designed to amplify the insert and to generate overlapping homologous sequences with the target vector, enabling seamless assembly using the Gibson Assembly method. Overhangs providing homology to the vector backbone are included at the 5′ ends of the primers, while the 3′ ends anneal to the template sequence. All primer sequences are listed in the 5′→3′ orientation.

| Fragments Gibson Assembly | 5' – 3' Sequence |
|---|---|
| pAAV6_AAVS1_FRAGMENT 1.FWD | CAGAGAGGGAGTGGCCAACTCCATCACTAGGGGTTCCTGCGGCCGCG ATTCGGGTCACCTCTCACTCC |
| pAAV6_AAVS1_FRAGMENT 1.REV | GGAGGAAGAGAAGAGGTCAGCTGTCCCTAGTGGCCCCAC |
| pAAV6_AAVS1_FRAGMENT 2.FWD | AGTGGGGCCACTAGGGACAGCTGACCTCTTCTCTTCCTCCCACAG |
| pAAV6_AAVS1_FRAGMENT 2.REV | GCCTGCAGGAATTGGTCGGGCCATAGAGCCCACCGCATCC |
| pAAV6_AAVS1_FRAGMENT 3.FWD | GGATGCGGTGGGCTCTATGGCCCGACCAATTC |
| pAAV6_AAVS1_FRAGMENT 3.REV | TTCATGGCGGGCATGGTGGCAGCTTCTCGAGTTCCAGGCAAACAG |
| pAAV6_AAVS1_FRAGMENT 4.FWD | TGCCTGGAACTCGAGAAGCTGCCACCATGCCCGCC |
| pAAV6_AAVS1_FRAGMENT 4.REV | AATGTATCTTATCATGTCTGCTAGACTCGAGATCTGGCGAAG |
| pAAV6_AAVS1_FRAGMENT 5.FWD | TCGCCAGATCTCGAGTCTAGCAGACATGATAAGATACATTGATGAGTTTG GACA |
| pAAV6_AAVS1_FRAGMENT 5.REV | GGGCTTTTCTGTCACCAATCAAAAAACCTCCCACACCTCCC |
| pAAV6_AAVS1_FRAGMENT 6.FWD | GGAGGTGTGGGAGGTTTTTTGATTGGTGACAGAAAAGCCCCC |
| pAAV6_AAVS1_FRAGMENT 6.REV | CAGAGAGGGAGTGGCCAACTCCATCACTAGGGGTTCCTGCGGCCGCA GACAGCCGCGTCAGAG |

**Table S2. PCR primers used for in–out PCR genotyping to confirm precise CD43 reporter knock-in.** Primer pairs were designed to verify correct genomic integration of the CD43-GFP donor template into the AAVS1 locus by in–out PCR genotyping. One primer anneals outside the homology arm in the genomic locus, while the second primer anneals within the inserted sequence, allowing amplification only when correct integration has occurred. Primer sequences are provided in the 5′→3′ orientation.

| **In-out PCR genotyping primers** | 5' – 3' Sequence |
|---|---|
| 5'_FWD | GATGCTCTTTCCGGAGCACT |
| 5'_REV | GATTCTCCTCCACGTCACCG |
| 3'_FWD | ACCGACAAGATCATCCGCAG |
| 3'_REV | GGGTGGCTACTGGCCTTATC |
